# Toxicological impacts and likely protein targets of bisphenol A in *Paramecium caudatum*

**DOI:** 10.1101/2021.06.24.449746

**Authors:** Marcus V. X. Senra, Ana Lúcia Fonseca

## Abstract

Bisphenol A (BPA) is a chemical agent widely used in plastic production and a well-known ubiquitous endocrine disruptor, frequently associated with a series of reproductive, developmental, and transgenerational impacts over wildlife, livestocks, and humans. Although widely studied, toxicological data on the effects of BPA are mostly restricted to mammalian models, remaining largely underexplored for other groups of organisms such as protists, which represents a considerable proportion of eukaryotic diversity. Here, we used acute end-point toxicological assay to evaluate the impacts of BPA over the survival of the cosmopolitan *Paramecium caudatum*; and a proteome-wide inverted virtual-screening (IVS) to predict the most likely *P. caudatum* proteins and pathways affected by BPA. This xenobiotic exerts a time-dependent effect over *P. caudatum* survival, which may be a consequence of impairments to multiple core cellular functions. We discuss the potential use of this ciliate as a biosensor for environmental BPA and as a new model organism to study the general impacts of this plasticizer agent over Eukaryotes. Finally, our data stress the relevance of bioinformatic methods to leverage the current knowledge on the molecular impacts of environmental contaminants over a diversity of biological systems.

## Introduction

The widespread contamination of aquatic ecosystems with a myriad of synthetic toxic compounds represents a significant threat to biodiversity (Karaouzas et al. 2018) and human safety (Wang et al. 2018), and can also intensify water shortage processes, through contamination of water supplies, all around the globe (Ma et al. 2020). Among these pollutants, bisphenol A (BPA), an organic synthetic compound largely employed in polycarbonate plastics and epoxy resins production has been attracting considerable attention due to its environmental ubiquity, consequence of its massive global demand, which is estimated in 3 million tons per year (Allard and Colaiácovo 2011), and because of its endocrine disruptor activity, frequently associated with a series of reproductive and developmental impairments over a variety of invertebrates and vertebrates (Flint et al. 2012), including humans (Abraham and Chakraborty 2019).

The impacts of BPA over exposed cells are broad (Aghajanpour-Mir et al. 2016; Can et al. 2005; Dumitrascu et al. 2020; Kundakovic and Champagne 2011) and are associated with impairments to a wide range of cellular pathways, among which the best studied protein targets are the nuclear hormone receptors, i.e. androgen, estrogen, and thyroid receptors (Usman and Ahmad 2016). In fact, according to Li and collaborators (Li et al. 2015), BPA can activate human estrogen receptor α (hERα), human estrogen-related receptor γ (hERRγ), and human peroxisome proliferator activated receptor γ (hPPARγ), by mimicking the structure of their natural agonist and binding to their corresponding site. Other cellular influences involves induction of oxidative stress (Yoon et al. 2014), apoptosis (Guo et al. 2017), and transgenerational epigenetic alterations through DNA methylation and histone acetylation pattern changes (Yoon et al. 2014) among many others.

Although extensive toxicological data are currently available, most of what is known about molecular impacts of BPA is restricted to metazoans and particularly, to mammalian cells (Flint et al. 2012). Therefore, the proteins targeted by BPA within unicellular microeukaryotes, organisms that are greatly diverse and crucial for the energy/carbon flow along trophic chains (Weisse 2017), remain largely underexplored.

Protein-ligand interactions can be determined through different *in vitro* approaches, such as crystallography (Delfosse et al. 2012), nuclear magnetic resonance (Yang et al. 2017), and competitive assays (Yang et al. 2016). However, these methods are extremely time-consuming, demand significant resources and their use in high-throughput analyses are greatly limited. Thus, in recent years, *in silico* approaches, such as molecular docking-based simulations, which could be used in both small (Li et al. 2015) and large scales (Xu et al. 2013) have been regarded as reliable alternative methods to overcome such limitations.

Among these docking-based approaches, the most widely used for proteome-wide target identification analysis is the inverted virtual screening (IVS), sometimes referred to as reverse virtual screening (Chen and Zhi 2001). In brief, this method starts with a searching algorithm to generate several possible binding modes between a given ligand and structures within a defined protein database, and then, an energy function is called to rank these predicted poses according to their free energy (Xu et al. 2018). Finally, proteins with the strongest binding energies are typically considered as the most likely targets for the desired ligand. In these terms, IVS could greatly boost our current understanding on the molecular mechanism of actions of a variety of environmental contaminants over model and non-model organisms with sequenced genomes.

Here, we used an acute end-point toxicological assay to investigate the detrimental impacts of BPA over the survival of an unicellular microeukaryote, the brackishwater and cosmopolitan ciliate (Ciliophora), *Paramecium caudatum*. Next, after predicting 3D protein structures based on the proteome of this species, we used IVS to identify likely protein targets of BPA, shedding some more light into the molecular bases underlying the effects of this xenobiotics over biological systems.

## Material and Methods

### Paramecium in vitro culture and acute toxicological assay

*In vitro* cultures of *Paramecium caudatum* strain JP1 (Boas et al. 2020) were maintained in petri dishes filled with 10 mL mineral water supplemented with macerated rice grains (Foissner et al. 2002). Stocks were kept at 23.0 °C and refreshed every week. Acute end-point toxicological assays were performed in 48-well polystyrene plates. Where, to each well 10 log-phase specimens were incubated for 2, 4, and 8h in the presence of 0.0001, 0,001, 0,01, 0,1, 1, 10, 100, or 1000 µM of BPA in mineral water at 23.0 °C. BPA stock solutions (Sigma-Aldrich, purity of ≥99%) were prepared to a concentration of 1 M in 50% DMSO (dimethyl sulfoxide) and then, serially diluted in mineral water to reach desirable concentrations. For this assay, a total of 5 experiments were performed, each one consisting in 5 replicates of each condition per time point. Mineral water and serial dilutions of 50% DMSO were used as negative controls. Dead ciliates were counted using a stereo microscope and mean lethal concentrations (LC50) were estimated for each time point using *Probit (Bliss 1935; Bliss 1934a; Bliss 1934b)*.

### Genome-wide spatial protein structure predictions

Since the genome sequence of *P. caudatum* strain JP1 is currently unavailable, we based our analyses on the reference genome of this species, *P. caudatum* strain 43c3d. Accordingly, its proteome (v.2), consisting of 18,673 predicted proteins, were retrieved from ParameciumDB (Arnaiz et al. 2020) and protein 3D structures were calculated using two homology-based modeling programs: *Modeller* v9.19 (Webb and Sali 2016) and *<u>P</u>rotein <u>H</u>omology/analog<u>Y R</u>ecognition <u>E</u>ngine* v.2 (*Phyre2*) (Kelley et al. 2015). For *Modeller*, best templates were selected using Blastp (Camacho et al. 2009) against proteins with defined 3D structures available from Protein Data Bank (PDB) (Berman et al. 2000). To ensure the quality of the models only proteins with sequence identities of >30% and coverage of >75% to the selected best template were included in the analysis. Then, 30 models for each *P. caudatum* protein were generated and the one with the lowest DOPE (Discrete Optimized Protein Structure) score, was selected for further analyses. Although *Phyre2* also applied a knowledge-based approach to generate models from the desired protein, it uses Hidden Markov Monte-Carlo (HMM) profile searches against PDB for best template selection, which can be more efficient to detect and to align distant homologs (Söding 2005), theoretically enhancing the quality of the final models. Again, the same sequence identity and coverage cutoff values used for Modeller were applied here to limit the universe of modeled proteins to the highest confidence possible. Next, model quality was accessed using *MolProbity (Chen et al. 2010)* and only the ones with ≥90% of amino acid residues within favorable regions of the Ramachandran plot were selected. In this last filtering step, when there were duplicate protein structures (modeled by both *Modeller* and *Phyre2*), the one with the highest number of amino acid residues within favorable regions of the Ramachandran plot were selected to feed the IVS assay.

### Inverse virtual screening (IVS)

Candidate protein targets for BPA within the P. *caudatum* modelled proteome were investigated using an IVS approach. The rationale here was based on the assumption that strong binding energies should indicate the most likely true associations. To start, all proteins were minimized in vacuum using *Gromacs* v5 (Abraham et al. 2015), applying OPLS-AA force field (Jorgensen et al. 1996) with 50.000 steps of steepest *descent* to resolve clashes and torsions. The search space, within each model, was defined by a grid enclosing the entire protein (blind docking) using *Obabel (O’Boyle et al. 2011)*. BPA 3D structure was directly retrieved from PubChem (https://pubchem.ncbi.nlm.nih.gov/), and we used *Obabel (O’Boyle et al. 2011)* to desalt, add polar hydrogens, and minimize (steepest descent) to resolve clashes and torsions. Then, an in-house python script was used to run *AutoDock Vina (Trott and Olson 2010)* and to compile the results. Afterwards, proteins with the lowest (strongest) binding energies to BPA, from the first percentile of frequency distribution, were selected as the most likely *P. caudatum* targets for this compound. Structure visualizations and image manipulations were done using *Chimera (Pettersen et al. 2004)*, and 2D diagrams of polar contacts between targeted proteins and BPA were done using *PoseView*, as implemented in *Protein Plus (Fährrolfes et al. 2017)*.

### Functional characterization of potential protein targets for BPA

Putative protein targets for BPA were characterized in terms of closest homologs, using *blastp* against SwissProt database (Bateman 2019); conserved domains searches, using *InterProScan (Jones et al. 2014), Prosite (de Castro et al. 2006)*, and *CD-search (Marchler-Bauer and Bryant 2004)*; subcellular location, using *Deeploc (Almagro Armenteros et al. 2017)*; pathways prediction, using *KEGG automatic annotation (Moriya et al. 2007)*; and cofactor and binding site predictions, using *Cofactor (Zhang et al. 2017)*.

### Functional enrichment analysis

To reconstruct the protein-protein interaction (PPI) network of the 34 putative protein targets for BPA, a functional enrichment analysis was performed querying the amino acid sequences of these proteins against the STRING database (Szklarczyk et al. 2019). All settings were in default mode with exception of the minimum required interaction score, which was changed to the highest confidence level (0.90) and max number of interactors, set to 100. PPI network was analysed and manipulated using *Cytoscape (Shannon et al. 2003)*.

## Results

To evaluate the impacts of BPA over the freshwater unicellular microeukaryote *Paramecium caudatum* strain JP1, acute endpoint toxicological assays were performed for 2, 4, and 8h and data are summarized in Table 1. As expected, BPA produced a time-dependent effect on ciliate’s survival, with the lowest LC_50_ value (7.42 µM or 15.28 mg/L) observed past 8h of exposure, indicating this widespread plasticizer agent, may also represent a potential environmental threat to this protist.

**Table 1.** Mean lethal concentration (LC_50_) of BPA over *P. caudatum* along exposures of 2, 4, and 8h.

| Exposure | $LC_{50}$ ($\mu M$) | Standard deviation (±) |
| --- | --- | --- |
| 2h | 43,641 | 2,839 |
| 4h | 16,727 | 1,586 |
| 8h | 7,415 | 2,048 |

We next asked what would be the molecular bases underlying this impact to *P. caudatum* survival rate and to address this issue, we used an inverted virtual screening (IVS) approach to identify the most likely *P. caudatum* protein targets for this plasticizer agent. The first step for this analysis was to model 3D structures of *P. caudatum* proteome based on data from the reference genome of this species (*P. caudatum* strain 43c3d), using two homology-based algorithms (*Modeller* and *Phyre2*). After successive rounds of filtering steps, 852 high-quality protein models were generated (4.5% of the predicted proteome size) (Table 2 and in Table S1), including members of 64 distinct cellular pathways, such as metabolic functions, ribosome-related enzymes, exosome membrane trafficking systems, and chromosome-associated proteins (Table S2).

**Table 2.**
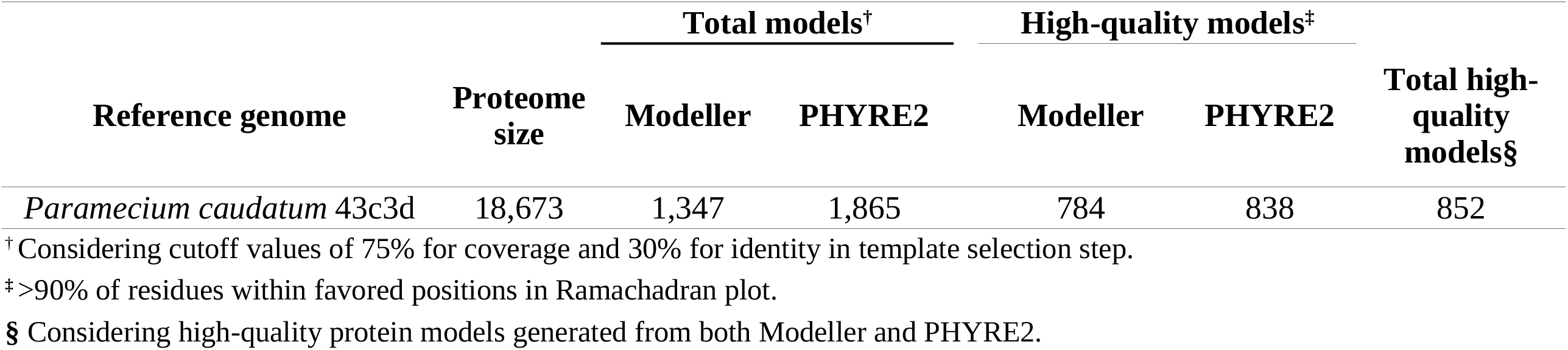
Summary of the *P. caudatum* proteome modelling process.

| Reference genome | Proteome size | Total models<sup>†</sup> |  | High-quality models<sup>‡</sup> |  | Total high-quality models<sup>§</sup> |
| --- | --- | --- | --- | --- | --- | --- |
|  |  | Modeller | PHYRE2 | Modeller | PHYRE2 |  |
| *Paramecium caudatum* 43c3d | 18,673 | 1,347 | 1,865 | 784 | 838 | 852 |

<sup>†</sup> Considering cutoff values of 75% for coverage and 30% for identity in template selection step.
<sup>‡</sup> >90% of residues within favored positions in Ramachadran plot.
<sup>§</sup> Considering high-quality protein models generated from both Modeller and PHYRE2.

Binding energies between these high-quality proteins and BPA were calculated using IVS and data are presented in Table S1. Such values varied considerably, ranging from strong values, such as -9.1 kcal/mol estimated for BPA and P00070322:Alkyl dihydroxyacetone phosphate synthase, to weak values, such as -3.8 kcal/mol between BPA and P00300095:RIO kinase. To improve the success rate of our predictions - correctly discriminate unlikely/spurious interactions from true associations - only proteins with the strongest binding energies to BPA, within the first percentile of the frequency distribution (<-7.9 kcal/mol), were considered for further investigations (Figure 1).

**Fig 1.**
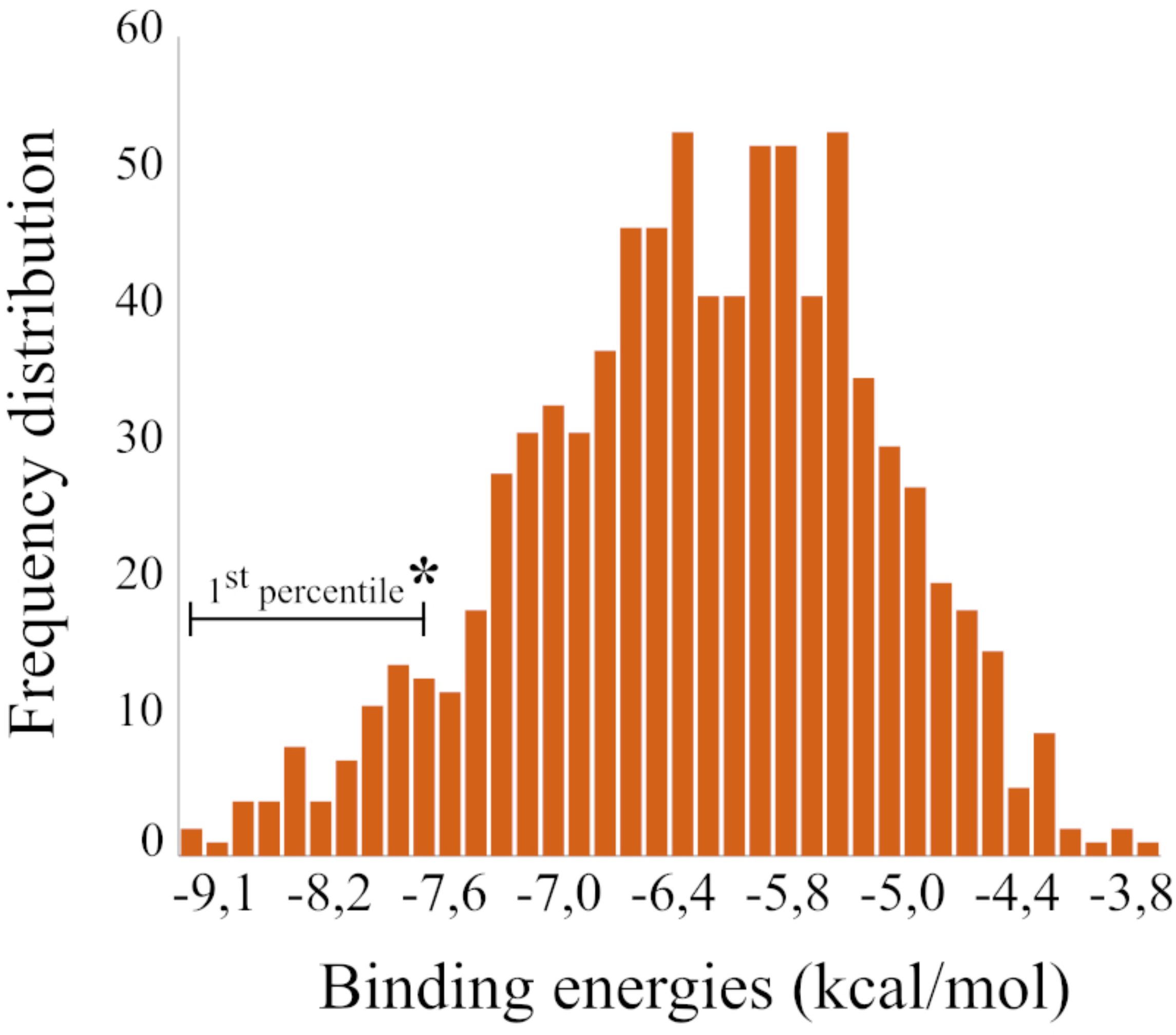
Frequency distribution of binding energies between BPA and 852 high-quality proteins modelled from the predicted proteome of *P. caudatum*. Third-four proteins with the strongest binding energies, indicated in the first percentile of the distributions were selected as likely targets for BPA

Thirty four proteins were identified within this group and were functionally annotated (Table 3 and Table S3). These proteins were predicted to different subcellular compartments, where they play crucial biological roles, such as DNA replication and cell cycle control, transcription and RNA splicing, protein transport and degradation, cell signaling, and lipid production among others. Many of these enzymes were assigned to one or more (KEGG) pathways (Table S3), and 10/34 can also interact with enzymes from distinct pathways, such as ribosome biogenesis, porphyrin metabolism, folate biosynthesis, and mismatch and nucleotide excision repair systems (Figure 2), as revealed by reconstructing the protein-protein interaction network of these 34 candidate protein targets for BPA, using data available from STRING database, suggesting that BPA may have a broad influence over *P. caudatum*.

**Table 3.** Functional annotation, binding modes, and binding energies of 34 putative protein targets of BPA.

| Protein ID<sup>†</sup> | Gene product | Biological process | Binding energies (mol/kcal) | Hydrogen bonds to BPA |
| --- | --- | --- | --- | --- |
| P00070270 | ABC transporter B family member 11 | Transmembrane transport (GO:0055085) | -8,3 | ARG 913/2HH2; ASN 71/OD1 |
| **P00300149** | **Protein kinase dsk1** | **Cell cycle (GO:0007049)** | **-8,7** | **GLU 196/OE2; GLU 196/N; ASN 134/1HD2; GLU 134/HN** |
| **P00090254** | **Phenylalanine-4-hydroxylase** | **Redoxi process (GO:0055114)** | **-8,6** | **TYR 365/OH** |
| P00030265 | Casein kinase I isoform epsilon | Regulation of G2/M transition (GO:0010389) | -8,4 | ASP 120/OD1 |
| **P00150104** | **Extracellular signal-regulated kinase 1** | **Activation of MAPK activity (GO:0000187)** | **-8,4** | **LYS 79/NZ** |
| P00210034 | Probable glutamine--tRNA ligase | Glutaminyl-tRNA aminoacylation (GO:0006425) | -8,4 | ARG 441/2HH1; GLU 436/OE2; LYS 209/O |
| **P00210141** | **Diphthine methyl ester synthase** | **Diphthamide biosynthesis (GO:0017183)** | **-8,3** | **CYS 91/O** |
| P00710081 | UDP-glucose 4-epimerase | Galactose metabolism (GO:0006012) | -8,3 | ARG 233/2HH2; ARG 233/2HH1 |
| P00100308 | Thymidylate kinase | dTDP biosynthesis (GO:0006233) | -8,2 | ASN 79/ND2 |
| P00420197 | Calmodulin | Calcium-mediated signaling (GO:0019722) | -8,2 | LYS 110/NZ |
| P00470113 | Extracellular signal-regulated kinase 1 | Protein phosphorylation (GO:0006468) | -8,2 | SER 192/HG; ASN 201/O; VAL 202/O |
| P00040164 | Dual specificity protein phosphatase CDC14A | Cell division (GO:0051301) | -8,1 | GLY 182/HN; ASP 164/O |
| P00870013 | Geranylgeranyl pyrophosphate synthase penG | Isoprenoid biosynthesis (GO:0008299) | -8,1 | ASP 46/OD1 |
| P00450170 | ABC transporter E family member 2 | Positive regulation of translation (GO:0045727) | -8 | GLN 171/O; SER 245/HG |
| P00300125 | 26S proteasome non-ATPase regulatory subunit 1 homolog B | Proteasome assembly (GO:0043248) | -8,4 | GLU 226/O; LYS 275/HZ2 |
| P00340021 | Calcium-transporting ATPase 2 | Calcium ion transport (GO:0006816) | -8,2 | PRO 398/O |
| **P00220032** | **Leucine aminopeptidase 2** | **Protein catabolism (GO:0030163)** | **-8** | **PRO 324/O; LYS 613/HN** |
| P00530165 | Calcium-transporting ATPase | Calcium ion transport (GO:1903515) | -8 | THR 236/HG1 |
| P00430133 | Cathepsin D | Proteolysis (GO:0006508) | -8,1 | PHE 106/O |
| **P00680119** | **Ras-related protein Rab-8B** | **Protein export (GO:0009306)** | **-8,5** | **SER 157/HG; LYS 159/HN; ALA 158/HN** |
| **P00260129** | **Ras-related protein Rab6** | **Protein transport (GO:0015031)** | **-8** | **SER 155/HG; LYS 157/HN; ALA 156/HN** |
| **P00700035** | **Ras-related protein ORAB-1** | **Protein transport (GO:0015031)** | **-8** | **LYS 155/HN; ALA 154/HN** |
| **P00680028** | **Phenylalanine-4-hydroxylase** | **Redoxi process (GO:0055114)** | **-8,9** | **GLU 273/OE1** |
| P00530125 | Peptidyl-prolyl cis-trans isomerase CYP19-2 | Protein folding (GO:0006457) | -8,7 | ASP 40/OD2; LYS 111/HN |
| P00620128 | Phenylalanine--tRNA ligase | Phenylalanyl-tRNA aminoacylation (GO:0006432) | -8,2 | PHE 197/O |
| P00800061 | Probable succinyl-CoA:3-ketoacid coenzyme A transferase | Cellular ketone body metabolism (GO:0046950) | -8,2 | ASN 303/2HD2 |
| **P00930049** | **Guanylate kinase 1** | **Purine nucleotide metabolism (GO:0006163)** | **-8** | **GLU 110/OE2** |
| P00370082 | DNA replication licensing factor mcm7 | DNA replication initiation (GO:0006270) | -8,2 | SER 275/HN; ASP 274/HN; LEU 142/O |
| **P00950041** | **Serine/threonine-protein kinase AFC3** | **Protein autophosphorylation (GO:0046777)** | **-8,6** | **LEU 127/HN** |
| **P00040322** | **Dual specificity protein kinase CLK4** | **Regulation of RNA splicing (GO:0043484)** | **-8,3** | **LEU 119/HN; SER 42/HG** |
| P00150260 | Cyclin-dependent kinase 10 | Regulation of mitosis (GO:0007346) | -8,1 | THR 191/O; VAL 190/O; GLU 221/OE2; LYS 155/HZ2 |
| P00300132 | DNA-directed RNA polymerase I subunit rpa1 | Transcription (GO:0006351) | -8 | TYR 369/HH |
| **P00100039** | **Peroxisomal acyl-coenzyme A oxidase 1** | **Fatty acid metabolism (GO:0006631)** | **-8,2** | **TYR 128/O; GLN 130/HN; THR 131/HN** |
| P00070322 | Alkyl-DHAP synthase | Lipid biosynthesis (GO:0008610) | -9,1 | ARG358/HH2; ARG 505/1HH2; ARG 505/HE |

Proteins in bold represent the ones in which BPA may occupy functionally important sites (see Table 4).
<sup>†</sup>According to original proteome annotation.

**Fig 2.**
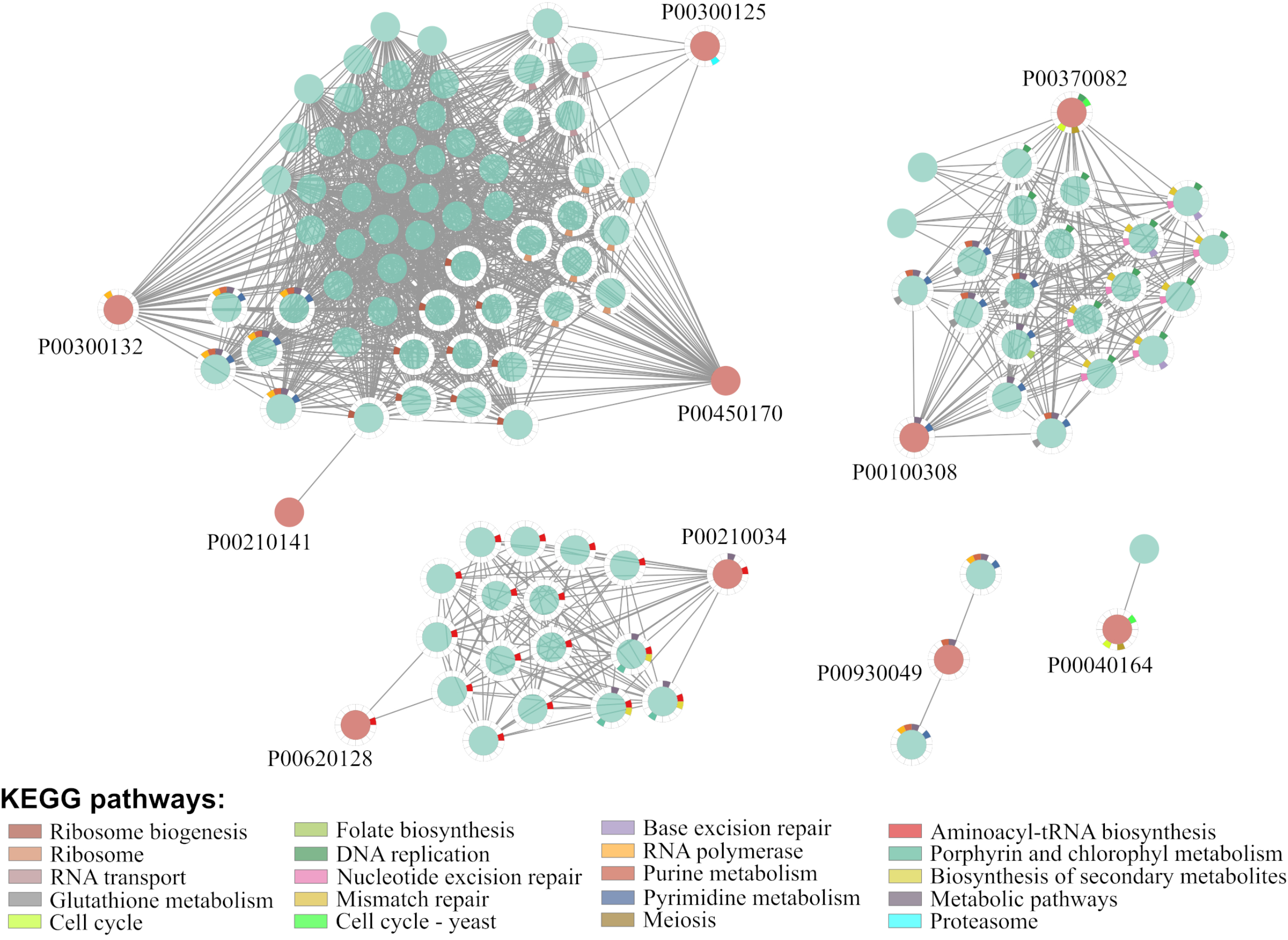
Protein-protein interaction (PPI) network of likely protein targets for BPA in *P. caudatum* based on data available from STRING database. Only candidate protein targets that interact with proteins from other pathways are presented. Circles in red represent the candidate protein targets, while in green are proteins retrieved during functional enrichment analysis. When known, the pathways to which each protein belongs are presented as color patterns displayed around the respective circle. P00040164 interacts with a kinase domain-containing protein (UNIPROT:A2DMG0) from *T. vaginalis* not allocated to any pathway

Among this group of proteins with strong binding energies to BPA (Table 3), 13/34 may interact with BPA within functionally important sites for cofactors (i.e. GDP/GTP), coenzymes (i.e. FAD) and inhibitors (i.e. staurosporine) (Table 4 and Figure S1), stressing that the predicted interactions between BPA and these enzymes may really produce biological effects.

**Table4.**
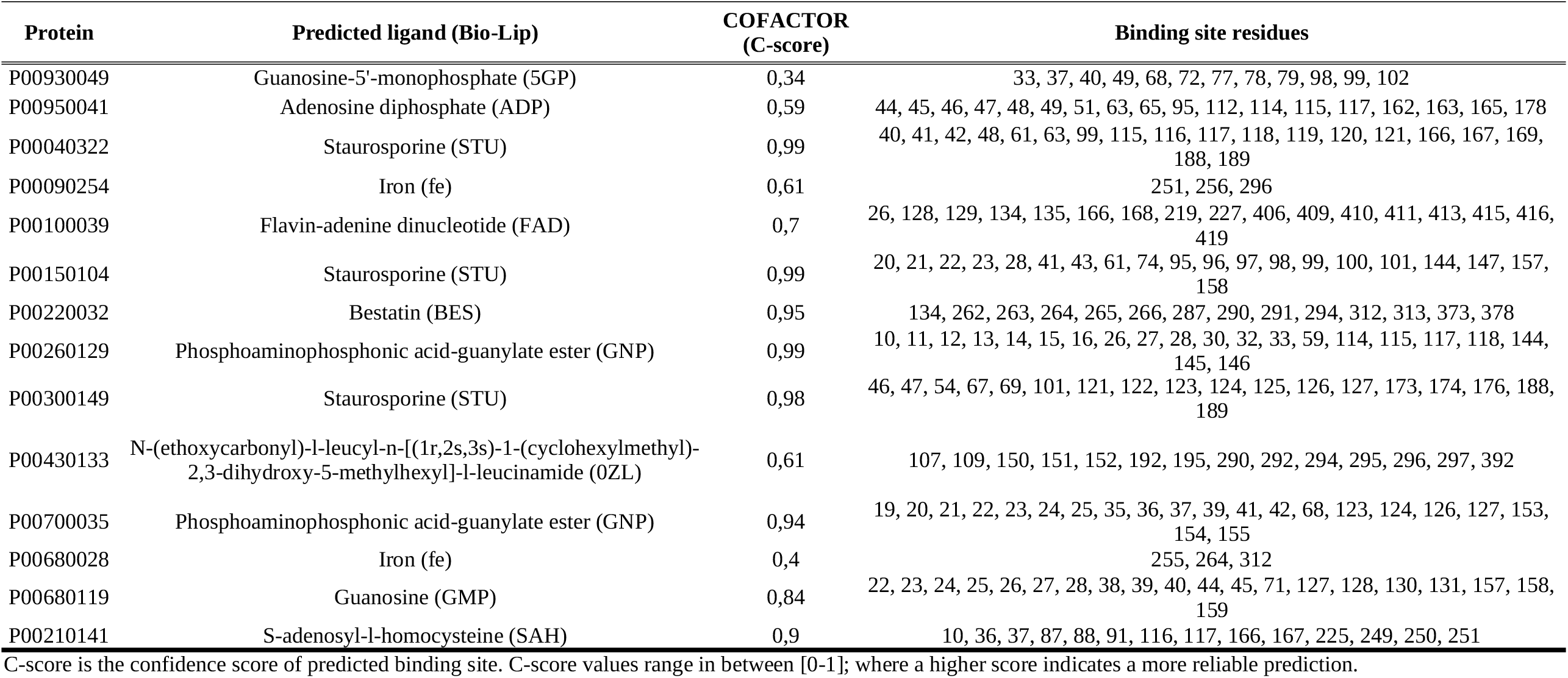
*P. caudatum* proteins that may be targeted by BPA within functionaly important sites, as predicted in analyses using *Cofactor*.

## Discussion

The freshwater ciliate *Paramecium caudatum* was firstly described by Ehrenberg in 1838 and, since then, mostly because of its worldwide distribution, simple taxonomy and *in vitro* maintenance, small body sizes, fast generation rates, and availability of genetic manipulation tools, *P. caudatum* have been serving as models in a vast array of disciplines, including cell biology (Sabaneyeva et al. 2009), climate change (Krenek et al. 2012), ecology (Violle et al. 2010), evolution (Johri et al. 2017), genetics and genomics (McGrath et al. 2014), and also toxicology (Boas et al. 2020).

Here, we used a *Paramecium caudatum* strain JP1, isolated from an eutrophic urban stream in Brazil (Boas et al. 2020), to evaluate how acute exposures to bisphenol A impacts its survival. The obtained LC _50_ measurements (Table 1) are relatively lower than values previously reported using a different *P. caudatum* strain sampled from an unrelated geographic region (Miyoshi et al. 2003). This should be a consequence of differences in their genetic background or even within the experimental conditions. Nevertheless, both estimates are within the same order of magnitude typically observed for invertebrates and vertebrates (Mathieu-Denoncourt et al. 2016), indicating this ciliate species should be regarded as a good biosensor for the environmental impacts of BPA.

Although BPA concentration in superficial waters (0-56 g/L) (Corrales et al. 2015) may not induce acute toxicological effects over *P. caudatum* nor to other aquatic organisms, it is now a consensus that long-term exposures, even at these environmental concentrations, are sufficient to produce diverse reproductive and developmental effects over wildlife (Flint et al. 2012) and epigenetic modifications in mice (Bansal et al. 2019; Wolstenholme et al. 2013) and in zebrafish (Santangeli et al. 2019). Considering the above mentioned benefits of using *Paramecium* as model organisms, we would like to highlight the great potential of this ciliate to improve our comprehension on the chronic effects of BPA over Eukaryotes.

Broad impacts are expected in *P. caudatum* during exposures to BPA. According to our data BPA can target 34 proteins from distinct subcellular compartments and different cellular functions, such as basic metabolism, DNA replication, transcription, translation, protein transport, signaling systems, and cell cycle (as schematized in Figure 3); many of which (10/34) able to interact with proteins of unrelated pathways (Figure 2), indicating that the observed effects on the survival of this ciliate may result from direct and indirect interferences over multiple core cellular functions.

**Fig 3.**
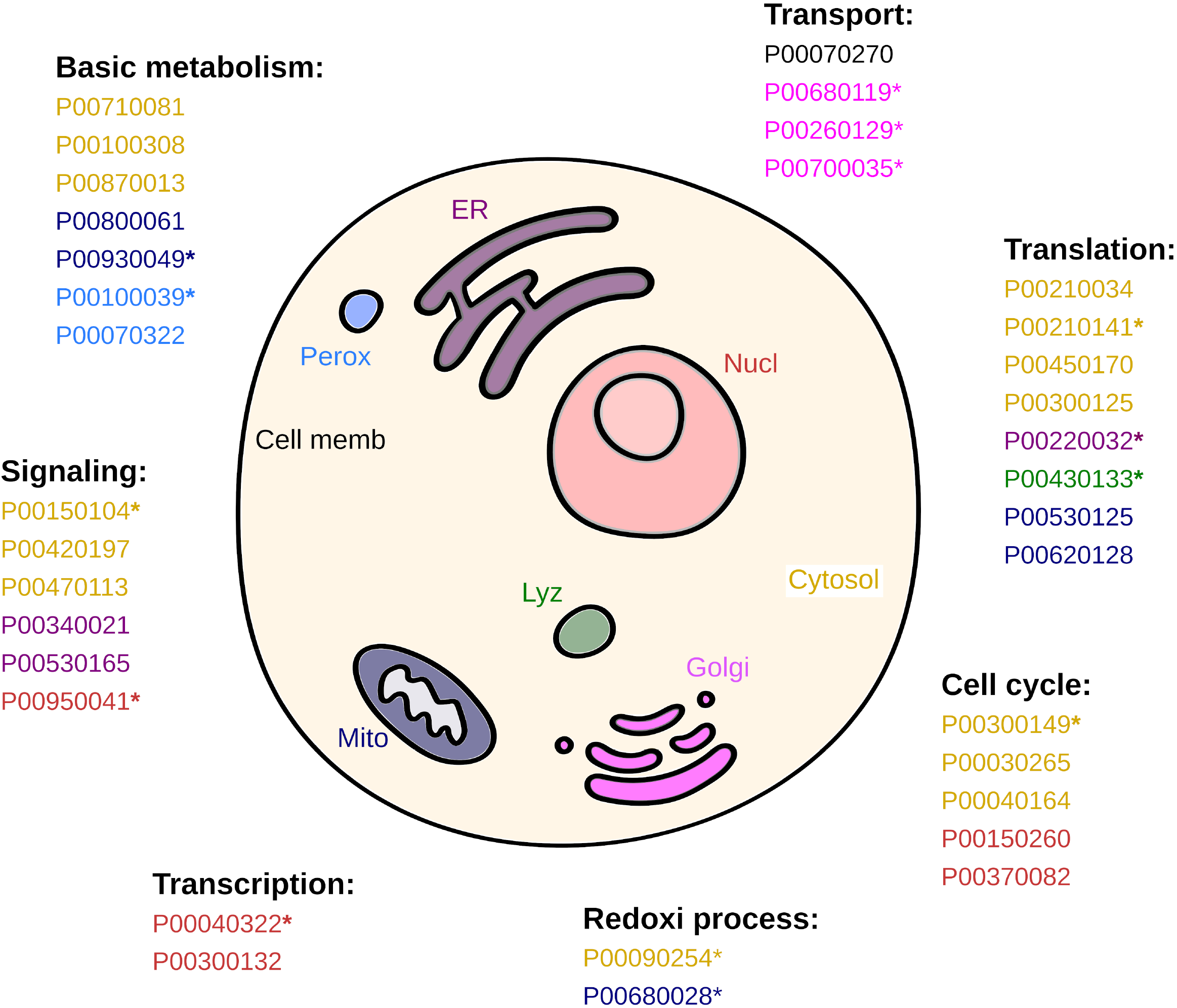
Schematic eukaryotic cell illustrating the putative protein targets for BPA in *P. caudatum* and their subcellular compartment and biological function. Protein codes are relative to Table 3. Proteins that may interact with BPA within the same site used by natural cofactors/coenzymes are marked (*)

Different lines of evidence stress that these predicted interactions may really occur in nature. For instance, binding energies measured for all 34 candidate targets are considerably lower than the typical accepted threshold value for stable protein-ligand associations (−6.0 kcal/mol) (Shityakov and Förster 2014) (Table 3); best binding poses calculated for many of these enzymes (14/34) are within conserved cofactor or coenzyme binding sites (Figure S1 and Table 4); and also, several likely affected pathways or biological functions (Table 3) have already been reported for other biological systems. On the other hand, whether such associations will cause enzyme activation, as described for small GTPase Ras proteins, after the binding of BPA to their GTP site (Schöpel et al. 2018; Schöpel et al. 2013) or inactivation, such for thyroid receptors, which occurs through the displacement of T3 hormone from the receptor (Moriyama et al. 2002), cannot be clearly accessed through bioinformatics and would require further investigations through biochemical approaches.

Different mechanisms can be used by BPA to cross plasmatic membranes and reach the cytosol of exposed cells. It can occur passively, after binding and inducing pore formation within plasmatic membranes (Chen et al. 2016), but also actively, through ATP-Binding Cassette (ABC) transporters (Mazur et al. 2012). Here, strong binding energy was measured between BPA and an ABCB-11 transporter (P00070270) within the plasmatic membrane of *P. caudatum* (Table 3). In fact, many ABCB transporters are involved in influx/efflux of a variety of metabolites in plants (Hwang et al. 2016); and interestingly, the best binding pose predicted in our analysis indicate that BPA may interact within amino acid residues facing the lumen side of the transmembrane channel (Figure S2), suggesting a plausible route of entrance toward *P. caudatum* cytoplasm.

Once in *P. caudatum* cytosol, BPA may transit to golgi apparatus, lysosome, mitochondria, nucleus, and endoplasmic reticulum, while potentially interfering with a wide range of cellular functions (Figure 3). Many of these interferences have already been described within other model organisms, such as extensive impacts over basic metabolic processes, including changes to the metabolism of galactose (Li et al. 2016), isoprenoid (Fic et al. 2015), lipid (Guan et al. 2019), ketone body (Meng et al. 2019), and purine (Li et al. 2018) (Table 3). In this sense, our data provide a possible molecular explanation for these observed phenotypic alterations. For instance, interferences to β-oxidation of fatty-acids within peroxisomes and purine metabolism within mitochondria may be a consequence of interaction of BPA within the binding sites flavin-adenine dinucleotide (FAD) within FAD-dependent acyl-CoA oxidase 1 (P00100039) and with the ATP site of guanylate kinase I (P00930049), the same used by staurosporine to inhibit the activity of a variety of kinases (Hirozane et al. 2019), respectively (Figure S1, and Table 4).

Besides the drastic transcriptomic (Yin et al. 2014), proteomic, and metabolomic (Huang et al. 2020) profile changes induced in organisms exposed to BPA, which may also occur in *P. caudatum* (although not evaluated here), our data suggest that general transcription and translation machineries are targets for BPA as well (Table 3). Concerning transcription, we predict impacts to rDNA transcription and RNA splicing, through interactions with RNA polymerase I (P00300132) (Table 3) and with dual specificity protein kinase CLK4 (P00040322), an enzyme that participates in the spliceosomal regulation through phosphorylating serine and arginine residues of members of this complex (Rosenthal et al. 2010), which may occur within its ATP binding site, the same region occupied by the alkaloid staurosporine (STU) to inhibit a variety of kinases (Zhou et al. 2006) (Figure S1, and Table 4).

Impacts over *P. caudatum* translation may be even more complex, involving impairments to tRNA biosynthesis (P00210034 and P00620128); ribosome export (P00450170), and protein degradation processes (P00300125, P00430133, and P00220032) (Table 3). Likewise, protein chain elongation and regular turnover may also be affected, since the high affinities found between BPA and diphthine methyl ester synthases (P00210141), within its coenzyme (S-adenosyl-L-homocysteine) site, an enzyme that participates in the unique post-translational modification of eukaryotic elongation factors (EF-2), resulting in the conversion of a histidine to diphthine residue, which is essential for its normal function (Zhu et al. 2010); and with leucine aminopeptidase 2 (Waditee-Sirisattha et al. 2011), within the same site used by bestatin (BES) to inhibit this enzyme activity (Stamper et al. 2004) (Figure S1, and Table S4).

Furthermore, serious effects to transport and signaling systems are also predicted. Three key regulators of intracellular membrane trafficking system (Gray et al. 2020), small GTPases Ras-related Rab proteins (P00680119, P00260129, and P00700035) are likely to interact with BPA through their GTP/GDP binding sites, which would result in impairments to vesicle and protein transport within this ciliate. With concerns to *P. caudatum*’ signaling systems, a likely target for BPA is the ubiquitous calcium signaling pathway (Ca^2+^/CaM/CaMKII) that regulates diverse calcium-dependent intracellular processes, including transcription, activation of regulatory enzymes, proliferation, and development, among others (Berridge et al. 2000). One of the major components of this pathway, calmodulin (CaM) (P00420197) (Table 3) is a likely target for BPA and; in fact, previous reports indicate that its interaction with BPA results in allosterically changes that reduce the ability to bind Ca2 ^+^ (Murayama et al. 2015), leading to male germ cell injuries in metazoans (Qian et al. 2015). In *paramecium*, motor functions (KINK et al. 1991) and trichocyst extrusion (Rauh and Nelson 1981) are controlled by calcium signaling pathway, suggesting that movement and cell defence mechanisms should also be impaired during exposures to BPA.

Impacts to *P. caudatum* cell cycle control is also predicted, in part a consequence of interactions of BPA with cytosolic extracellular signal-regulated kinase 1 (ERK) (P00150104 and P0004016) (Table 3, Figure S1, and Table 4). These enzymes are members of the conserved ERK pathway, which regulates a diversity of cellular functions, including cell entrance in mitosis and meiosis (Liu et al. 2004). Therefore, impairments to these enzymes’ normal functions would lead to G1/S cell arrestments, as previously reported in other organisms (Bilancio et al. 2017; Chu et al. 2018). Moreover, BPA would likely interfere within G2/M checkpoint of *P. caudatum*, as proposed to occur in mice (Liu et al. 2013), given the strong binding energies observed with cyclin-dependent kinase 14 (CDK14) (P00150260), a cell cycle regulator involved in G2/M transition (Wang et al. 2016); with DNA replication licensing factor mcm7 (P00370082), a participant of the MCM2-7 complex, which is essential for the initiation of DNA replication and elongation throughout Eukaryotes (Liang et al. 2017); and with protein kinase dsk1 (P00300149), a protein involved in G2/M progression in Yeast (Takeuchi and Yanagida 1993). Still, we believe that BPA may also influentiate normal cytokinesis and cause mitotic exit of *P. caudatum* by interacting with Dual specificity protein phosphatase CDC14A (P00040164), an key enzyme responsible for the down-regulation of mitotic Cdks (Gray et al. 2003) (Table 3).

Finally, BPA can also interact with two Phenylalanine-4-hydroxylases (EC:1.14.16.1) (P00090254 and P00680028), which are oxidoreductases involved in the conversion of L-phenylalanine to L-tyrosine. The configuration assumed by BPA, with one of its aromatic rings oriented toward the Fe_2_^3+^ center (FigureS3), resembles the binding mode of L-phenylalanine within the active site of these enzymes (Andreas Andersen et al. 2002). Therefore, we hypothesise that BPA could act either as an competitive inhibitor or as a substrate for these two oxidoreductases, which could have a role in detoxification processes of intracellular BPA.

## Conclusion

As a consequence of the enormous world-wide demand for plastic production and its recalcitrant properties, BPA can be considered a ubiquitous and persistent environmental contaminant, with great potential to cause acute and chronic damages to a variety of microeukaryotes, invertebrates, and vertebrates. Therefore, it should be closely monitored and more effective legislation and risk management programs should be established to mitigate its impacts. In this context, we showed that the brackishwater cosmopolitan ciliate *Paramecium caudatum* could be a good biosensor for environmental BPA and also an efficient model to better evaluate both acute and chronic toxicological impacts of this plasticizer agent.

The cellular impacts of BPA over *P. caudatum* are broad and may involve interferences to multiple conserved core functions. Many new molecular targets for BPA were uncovered here and since they are conserved throughout Eukaryotes, our predictions should be further evaluated in other unicellular and multicellular organisms. Moreover, we believe that the framework applied here, using molecular docking approaches, should be used more often to provide a better comprehension on how contaminants of emerging concern affect a wide variety of biological systems.

## Acknowledgements

The authors acknowledge the “Coordenação de Aperfeiçoamento de Pessoal de Nível Superior” (CAPES) for the post-doctoral fellowship (PNPD) (88882.317976/2019-01) conferred to MS.

## Declarations

### Availability of data and material

Data and material will be available at request.

### Code availability

Code will be available at request.

### Authors’ contributions

Marcus Senra: Conceptualization, methodology, investigation, and writing-original draft preparation: Ana Fonseca: Supervision, writing-reviewing, and editing.

## Figure captions

**Supplementary Fig S1** Three-dimensional structures of 14 candidate protein targets for BPA in *P. caudatum*, in which the predicted besting binding pose is located within regions used by the natural cofactor/coenzyme of the enzyme according to analysis using COFACTOR. Each candidate protein target is represented twice: In the top part is presented the best binding pose for BPA (in purple); and in the bottom part the binding site used by the cofactor/coenzyme (in red)

**Supplementary Fig S2** Three-dimensional structure of candidate protein targets P00070270, an ABC transporter B family member 11 of *P. caudatum*, which may be involved in the translocation of BPA from extracellular region to cytosol. The predicted best binding pose for BPA is represented in purple and occurs within the lumen-side of the transmembrane channel

**Supplementary Fig S3** Three-dimensional structures of 2 candidate protein targets for BPA in *P. caudatum*, which could be involved in detoxification processes, through oxidation, to protect cells from impacts caused by BPA. (A) The cytoplasmic P00090254 and (B) the mitochondrial P00680028 Phenylalanine-4-hydroxylases. Iron ions are represented as a golden sphere and BPA are in purple. The small boxes depict hydrophobic (green spindles) and polar (dotted lines) interactions between BPA and amino acid residues from the enzymes

