## Supplementary material for "Toxicological impacts and likely protein targets of bisphenol A in *Paramecium caudatum*": Supp. Material

**Table S1.** Annotation data on the *P. caudatum* 3D modelled proteins and their binding energies to BPA.

| ID | DESCRIPTION | CHROMOSOME | NT_START | NT_END | BINDING ENERGIES (kcal/mol) |
|---|---|---|---|---|---|
| PCAU.43c3d.1.P00010012 | Tryptophan synthase beta subunit-like PLP-dependent enzyme | scaffold_0001 | 23197 | 24853 | -7.4 |
| PCAU.43c3d.1.P00010015 | Ribosomal protein L32e | scaffold_0001 | 26373 | 26859 | -6.2 |
| PCAU.43c3d.1.P00010044 | Catalase, mono-functional, haem-containing | scaffold_0001 | 71821 | 73367 | -6.5 |
| PCAU.43c3d.1.P00010050 | Dihydroorotate dehydrogenase, class 1/ 2 | scaffold_0001 | 76614 | 79650 | -6.6 |
| PCAU.43c3d.1.P00010054 | Serine/threonine/dual specificity protein kinase, catalytic domain | scaffold_0001 | 83399 | 84653 | -6.7 |
| PCAU.43c3d.1.P00010070 | Peptidyl-prolyl cis-trans isomerase, FKBP-type | scaffold_0001 | 104909 | 105387 | -5.9 |
| PCAU.43c3d.1.P00010103 | V-ATPase proteolipid subunit C-like domain | scaffold_0001 | 168736 | 169346 | -5.6 |
| PCAU.43c3d.1.P00010112 | DNA-directed RNA polymerase, RBP11-like dimerisation domain | scaffold_0001 | 180310 | 181372 | -6.0 |
| PCAU.43c3d.1.P00010165 | Vacuolar (H+)-ATPase G subunit | scaffold_0001 | 252653 | 253112 | -5.6 |
| PCAU.43c3d.1.P00010176 | Coproporphyrinogen III oxidase, aerobic | scaffold_0001 | 262051 | 263168 | -6.7 |
| PCAU.43c3d.1.P00010205 | Metalloenzyme, LuxS/M16 peptidase-like | scaffold_0001 | 299060 | 301881 | -7.4 |
| PCAU.43c3d.1.P00010249 | P-loop containing nucleoside triphosphate hydrolase | scaffold_0001 | 378899 | 380283 | -6.6 |
| PCAU.43c3d.1.P00010267 | Translation Initiation factor eIF- 4e | scaffold_0001 | 406368 | 406891 | -5.5 |
| PCAU.43c3d.1.P00010325 | ATP:guanido phosphotransferase, catalytic domain | scaffold_0001 | 496004 | 497221 | -6.9 |
| PCAU.43c3d.1.P00010342 | Cyclophilin-like domain | scaffold_0001 | 524342 | 524955 | -5.7 |
| PCAU.43c3d.1.P00010366 | Ribosomal protein S26e | scaffold_0001 | 568059 | 568545 | -6.7 |
| PCAU.43c3d.1.P00010367 | EF-hand domain pair | scaffold_0001 | 568635 | 569075 | -6.2 |
| PCAU.43c3d.1.P00010434 | Nucleophile aminohydrolases, N-terminal | scaffold_0001 | 664238 | 665000 | -6.8 |
| PCAU.43c3d.1.P00010441 | ATP:guanido phosphotransferase, catalytic domain | scaffold_0001 | 671664 | 672981 | -6.4 |
| PCAU.43c3d.1.P00010442 | ATP:guanido phosphotransferase, catalytic domain | scaffold_0001 | 673343 | 674561 | -6.1 |
| PCAU.43c3d.1.P00010445 | Heat shock protein Hsp90 family | scaffold_0001 | 676202 | 678364 | -6.3 |
| PCAU.43c3d.1.P00010472 | Nucleotide-binding, alpha-beta plait | scaffold_0001 | 742728 | 743261 | -6.0 |
| PCAU.43c3d.1.P00010491 | DNA-directed RNA polymerase, RBP11-like dimerisation domain | scaffold_0001 | 776121 | 777138 | -6.7 |
| PCAU.43c3d.1.P00020002 | Metallo-dependent phosphatase-like | scaffold_0002 | 2096 | 3154 | -6.3 |
| PCAU.43c3d.1.P00020017 | MEMO1 family | scaffold_0002 | 22159 | 23173 | -6.2 |
| PCAU.43c3d.1.P00020018 | V-type ATPase, V0 complex, 116kDa subunit family | scaffold_0002 | 23337 | 26101 | -7.0 |
| PCAU.43c3d.1.P00020020 | ATPase, V0 complex, subunit d | scaffold_0002 | 28975 | 30278 | -7.8 |
| PCAU.43c3d.1.P00020032 | Plectin/S10, N-terminal | scaffold_0002 | 46215 | 46741 | -6.2 |
| PCAU.43c3d.1.P00020086 | small GTPase Rab1 family profile. | scaffold_0002 | 124395 | 124912 | -7.3 |
| PCAU.43c3d.1.P00020129 | Zinc-binding ribosomal protein | scaffold_0002 | 179779 | 180443 | -5.4 |
| PCAU.43c3d.1.P00020184 | ClpP/crotonase-like domain | scaffold_0002 | 273785 | 274707 | -6.1 |
| PCAU.43c3d.1.P00020185 | Nucleophile aminohydrolases, N-terminal | scaffold_0002 | 274672 | 275509 | -6.5 |
| PCAU.43c3d.1.P00020219 | Peptidase C1A, papain C-terminal | scaffold_0002 | 340060 | 341619 | -6.6 |
| PCAU.43c3d.1.P00020241 | P-loop containing nucleoside triphosphate hydrolase | scaffold_0002 | 386885 | 388139 | -6.2 |
| PCAU.43c3d.1.P00020266 | Ribosomal protein S5 domain 2-type fold, subgroup | scaffold_0002 | 452040 | 452580 | -5.5 |
| PCAU.43c3d.1.P00020276 | Ribosomal protein L24e-related | scaffold_0002 | 473300 | 473907 | -6.6 |
| PCAU.43c3d.1.P00020302 | Ribosomal protein L29e | scaffold_0002 | 520059 | 520369 | -5.8 |
| PCAU.43c3d.1.P00020306 | Ubiquitin-like | scaffold_0002 | 527452 | 528002 | -6.1 |
| PCAU.43c3d.1.P00020310 | P-loop containing nucleoside triphosphate hydrolase | scaffold_0002 | 531352 | 532168 | -6.9 |
| PCAU.43c3d.1.P00020311 | Triosephosphate isomerase | scaffold_0002 | 532158 | 532970 | -6.5 |
| PCAU.43c3d.1.P00020337 | Ribosomal protein S5/S7 | scaffold_0002 | 566450 | 567191 | -6.1 |
| PCAU.43c3d.1.P00020367 | Rossmann-like alpha/beta/alpha sandwich fold | scaffold_0002 | 616707 | 617803 | -6.5 |
| PCAU.43c3d.1.P00020370 | P-loop containing nucleoside triphosphate hydrolase | scaffold_0002 | 622249 | 623018 | -6.4 |
| PCAU.43c3d.1.P00030023 | Protein kinase-like domain | scaffold_0003 | 37821 | 38943 | -6.2 |
| PCAU.43c3d.1.P00030149 | Ribosomal protein L11/L12 | scaffold_0003 | 221344 | 221912 | -6.3 |
| PCAU.43c3d.1.P00030183 | small GTPase Rab1 family profile. | scaffold_0003 | 269552 | 270222 | -6.1 |
| PCAU.43c3d.1.P00030230 | Ribosomal protein L2 | scaffold_0003 | 334811 | 335773 | -7.3 |
| PCAU.43c3d.1.P00030248 | Thioredoxin-like fold | scaffold_0003 | 370296 | 371093 | -6.0 |
| PCAU.43c3d.1.P00030253 | Pyruvate/Phosphoenolpyruvate kinase-like domain | scaffold_0003 | 378174 | 379185 | -6.4 |
| PCAU.43c3d.1.P00030265 | Protein kinase-like domain | scaffold_0003 | 392077 | 393317 | -8.4 |
| PCAU.43c3d.1.P00030294 | Tubulin | scaffold_0003 | 438362 | 439629 | -6.8 |
| PCAU.43c3d.1.P00030304 | Glutathione S-transferase, C-terminal-like | scaffold_0003 | 451438 | 452186 | -5.9 |
| PCAU.43c3d.1.P00030310 | Malate dehydrogenase, type 2 | scaffold_0003 | 457945 | 459013 | -7.4 |
| PCAU.43c3d.1.P00030338 | Ribosomal protein L15e | scaffold_0003 | 502989 | 503719 | -5.8 |
| PCAU.43c3d.1.P00030345 | Ribosomal protein S15 | scaffold_0003 | 510311 | 510915 | -6.3 |
| PCAU.43c3d.1.P00030361 | Nucleotide-diphospho-sugar transferases | scaffold_0003 | 524711 | 525492 | -7.3 |
| PCAU.43c3d.1.P00030375 | cAMP/cGMP-dependent protein kinase | scaffold_0003 | 542438 | 543602 | -5.7 |
| PCAU.43c3d.1.P00030386 | Ribosomal protein L34Ae | scaffold_0003 | 558978 | 559397 | -5.5 |
| PCAU.43c3d.1.P00030410 | DNA-directed RNA polymerase, RBP11-like dimerisation domain | scaffold_0003 | 597253 | 597642 | -5.8 |
| PCAU.43c3d.1.P00030421 | Small GTPase superfamily, ARF type | scaffold_0003 | 608142 | 608790 | -5.9 |
| PCAU.43c3d.1.P00040013 | NAD(P)-binding domain | scaffold_0004 | 18022 | 19503 | -7.0 |
| PCAU.43c3d.1.P00040043 | Ammonium transporter | scaffold_0004 | 64877 | 66436 | -6.4 |
| PCAU.43c3d.1.P00040075 | Tubulin | scaffold_0004 | 111938 | 113360 | -6.6 |
| PCAU.43c3d.1.P00040081 | Ribosomal protein L22e | scaffold_0004 | 118361 | 118815 | -5.8 |
| PCAU.43c3d.1.P00040164 | Tyrosine-protein phosphatase CDC14 | scaffold_0004 | 249073 | 250499 | -8.1 |
| PCAU.43c3d.1.P00040193 | RmlC-like jelly roll fold | scaffold_0004 | 293908 | 295104 | -6.1 |
| PCAU.43c3d.1.P00040264 | Cyclin-dependent kinase, regulatory subunit | scaffold_0004 | 406649 | 406953 | -5.1 |
| PCAU.43c3d.1.P00040322 | Protein kinase-like domain | scaffold_0004 | 496132 | 497256 | -8.3 |
| PCAU.43c3d.1.P00050046 | Phosphoribosyltransferase-like | scaffold_0005 | 68698 | 69266 | -5.7 |
| PCAU.43c3d.1.P00050099 | Ubiquitin-conjugating enzyme/RWD-like | scaffold_0005 | 151495 | 152131 | -6.0 |
| PCAU.43c3d.1.P00050122 | Glutathione S-transferase, C-terminal-like | scaffold_0005 | 184658 | 185392 | -7.4 |
| PCAU.43c3d.1.P00050123 | Ribosomal protein S30 | scaffold_0005 | 185373 | 185722 | -6.6 |
| PCAU.43c3d.1.P00050128 | Biotin/lipoate A/B protein ligase | scaffold_0005 | 190964 | 191992 | -7.6 |
| PCAU.43c3d.1.P00050150 | Alpha-D-phosphohexomutase, alpha/beta/alpha I/II/III | scaffold_0005 | 236856 | 238711 | -7.5 |
| PCAU.43c3d.1.P00050155 | P-loop containing nucleoside triphosphate hydrolase | scaffold_0005 | 248384 | 249926 | -7.3 |
| PCAU.43c3d.1.P00050208 | Peptidase M1, alanine aminopeptidase/leukotriene A4 hydrolase | scaffold_0005 | 340131 | 342182 | -7.6 |
| PCAU.43c3d.1.P00050211 | P-loop containing nucleoside triphosphate hydrolase | scaffold_0005 | 344515 | 346995 | -6.4 |
| PCAU.43c3d.1.P00050228 | Aconitase/2-methylisocitrate dehydratase | scaffold_0005 | 362142 | 364909 | -7.0 |
| PCAU.43c3d.1.P00050243 | P-loop containing nucleoside triphosphate hydrolase | scaffold_0005 | 386135 | 386857 | -6.1 |
| PCAU.43c3d.1.P00050246 | GNAT domain | scaffold_0005 | 389146 | 389709 | -6.9 |
| PCAU.43c3d.1.P00050250 | P-loop containing nucleoside triphosphate hydrolase | scaffold_0005 | 392574 | 394251 | -6.3 |
| PCAU.43c3d.1.P00050294 | Chaperonin Cpn60/TCP-1 | scaffold_0005 | 472540 | 474281 | -7.1 |
| PCAU.43c3d.1.P00050317 | Ribosomal protein L24e-related | scaffold_0005 | 500106 | 500725 | -6.5 |
| PCAU.43c3d.1.P00050320 | AAA+ ATPase domain | scaffold_0005 | 503245 | 505606 | -7.1 |

| | | | | | |
|---|---|---|---|---|---|
| PCAU.43c3d.1.P00050337 | Protein phosphatase 2C | scaffold_0005 | 528684 | 529629 | -6.2 |
| PCAU.43c3d.1.P00060007 | Ubiquitin-like | scaffold_0006 | 10590 | 11105 | -5.7 |
| PCAU.43c3d.1.P00060008 | Ubiquitin-like | scaffold_0006 | 11570 | 12080 | -5.7 |
| PCAU.43c3d.1.P00060009 | Ubiquitin-like | scaffold_0006 | 12089 | 12540 | -5.2 |
| PCAU.43c3d.1.P00060010 | Ubiquitin-like | scaffold_0006 | 14369 | 14860 | -5.4 |
| PCAU.43c3d.1.P00060011 | Ubiquitin-like | scaffold_0006 | 15437 | 16129 | -6.4 |
| PCAU.43c3d.1.P00060029 | Peptidase C12, ubiquitin carboxyl-terminal hydrolase | scaffold_0006 | 43998 | 44815 | -6.6 |
| PCAU.43c3d.1.P00060039 | P-loop containing nucleoside triphosphate hydrolase | scaffold_0006 | 60312 | 61613 | -6.0 |
| PCAU.43c3d.1.P00060041 | Thiamin diphosphate-binding fold | scaffold_0006 | 62370 | 63614 | -6.4 |
| PCAU.43c3d.1.P00060076 | Ribosomal protein L1, 2-layer alpha/beta-sandwich | scaffold_0006 | 109196 | 109964 | -6.3 |
| PCAU.43c3d.1.P00060077 | P-loop containing nucleoside triphosphate hydrolase | scaffold_0006 | 110191 | 112854 | -7.4 |
| PCAU.43c3d.1.P00060084 | DNA topoisomerase I, eukaryotic-type | scaffold_0006 | 119205 | 121047 | -6.9 |
| PCAU.43c3d.1.P00060103 | NADH:cytochrome b5 reductase (CBR) | scaffold_0006 | 160952 | 161959 | -6.9 |
| PCAU.43c3d.1.P00060111 | EF-hand domain pair | scaffold_0006 | 172009 | 172531 | -5.5 |
| PCAU.43c3d.1.P00060115 | Tubulin | scaffold_0006 | 178024 | 179283 | -6.8 |
| PCAU.43c3d.1.P00060120 | Tetratricopeptide-like helical domain | scaffold_0006 | 183651 | 185047 | -6.7 |
| PCAU.43c3d.1.P00060133 | Ribosomal protein L13 | scaffold_0006 | 201102 | 201802 | -6.5 |
| PCAU.43c3d.1.P00060147 | Ribosomal protein L44e | scaffold_0006 | 216816 | 217218 | -6.1 |
| PCAU.43c3d.1.P00060164 | Ribosomal protein L31e | scaffold_0006 | 238542 | 239005 | -6.1 |
| PCAU.43c3d.1.P00060173 | Copper homeostasis protein CutC | scaffold_0006 | 251481 | 251979 | -5.6 |
| PCAU.43c3d.1.P00060216 | Glycogen debranching enzyme | scaffold_0006 | 326118 | 330796 | -7.5 |
| PCAU.43c3d.1.P00060230 | Armadillo-type fold | scaffold_0006 | 353462 | 355407 | -7.3 |
| PCAU.43c3d.1.P00060343 | small GTPase Rab1 family profile. | scaffold_0006 | 528268 | 529015 | -7.4 |
| PCAU.43c3d.1.P00070009 | Pyruvate kinase | scaffold_0007 | 20914 | 22568 | -6.1 |
| PCAU.43c3d.1.P00070047 | Ribosomal protein S11 | scaffold_0007 | 73837 | 74458 | -6.0 |
| PCAU.43c3d.1.P00070048 | Ribosomal protein L19/L19e domain | scaffold_0007 | 74501 | 75170 | -5.3 |
| PCAU.43c3d.1.P00070057 | P-type ATPase, transmembrane domain | scaffold_0007 | 90433 | 93796 | -7.6 |
| PCAU.43c3d.1.P00070098 | Ribosomal protein L37e | scaffold_0007 | 158310 | 158683 | -5.3 |
| PCAU.43c3d.1.P00070127 | Glutamine synthetase, catalytic domain | scaffold_0007 | 196111 | 197392 | -6.1 |
| PCAU.43c3d.1.P00070147 | Autophagy protein Atg8 ubiquitin like | scaffold_0007 | 222957 | 223363 | -6.4 |
| PCAU.43c3d.1.P00070149 | small GTPase Rab1 family profile. | scaffold_0007 | 226906 | 227682 | -6.3 |
| PCAU.43c3d.1.P00070152 | EF-hand domain pair | scaffold_0007 | 229155 | 229707 | -5.7 |
| PCAU.43c3d.1.P00070175 | Urocanase | scaffold_0007 | 256544 | 258799 | -7.8 |
| PCAU.43c3d.1.P00070202 | Acyl-CoA dehydrogenase/oxidase C-terminal | scaffold_0007 | 304758 | 306871 | -7.1 |
| PCAU.43c3d.1.P00070233 | Protein kinase-like domain | scaffold_0007 | 341722 | 342755 | -7.9 |
| PCAU.43c3d.1.P00070236 | Chaperonin Cpn60/TCP-1 | scaffold_0007 | 344275 | 346204 | -7.8 |
| PCAU.43c3d.1.P00070238 | P-loop containing nucleoside triphosphate hydrolase | scaffold_0007 | 348547 | 349320 | -7.0 |
| PCAU.43c3d.1.P00070268 | Aminoacyl-tRNA synthetase, class II (D/K/N)-like | scaffold_0007 | 404503 | 406555 | -6.7 |
| PCAU.43c3d.1.P00070270 | ABC transporter, transmembrane domain | scaffold_0007 | 408333 | 412384 | -8.3 |
| PCAU.43c3d.1.P00070272 | P-type ATPase, transmembrane domain | scaffold_0007 | 413623 | 417104 | -7.3 |
| PCAU.43c3d.1.P00070293 | Heat shock protein Hsp90 family | scaffold_0007 | 445442 | 447645 | -6.9 |
| PCAU.43c3d.1.P00070294 | Ribosomal protein S7e | scaffold_0007 | 448029 | 448779 | -6.1 |
| PCAU.43c3d.1.P00070322 | Alkyldihydroxyacetonephosphate synthase | scaffold_0007 | 497260 | 499237 | -9.1 |
| PCAU.43c3d.1.P00080012 | Glycyl-tRNA synthetase/DNA polymerase subunit gamma-2 | scaffold_0008 | 11786 | 13824 | -7.2 |
| PCAU.43c3d.1.P00080040 | Ribosomal protein L35A | scaffold_0008 | 54476 | 54972 | -6.4 |
| PCAU.43c3d.1.P00080041 | Chaperonin Cpn60/TCP-1 | scaffold_0008 | 54980 | 56759 | -5.9 |
| PCAU.43c3d.1.P00080045 | small GTPase Rab1 family profile. | scaffold_0008 | 59776 | 60382 | -6.7 |
| PCAU.43c3d.1.P00080057 | Fatty acid desaturase, type 1, core | scaffold_0008 | 72323 | 73332 | -6.9 |
| PCAU.43c3d.1.P00080063 | 14-3-3 domain | scaffold_0008 | 83736 | 84709 | -5.9 |
| PCAU.43c3d.1.P00080115 | small GTPase Rab1 family profile. | scaffold_0008 | 180783 | 181495 | -7.2 |
| PCAU.43c3d.1.P00080127 | Protein kinase-like domain | scaffold_0008 | 195202 | 197807 | -6.9 |
| PCAU.43c3d.1.P00080133 | Cyclic nucleotide-binding domain | scaffold_0008 | 204010 | 205107 | -6.2 |
| PCAU.43c3d.1.P00080141 | Flavoprotein pyridine nucleotide cytochrome reductase | scaffold_0008 | 216008 | 218077 | -6.6 |
| PCAU.43c3d.1.P00080143 | Ribosomal protein S19e | scaffold_0008 | 218464 | 219027 | -6.6 |
| PCAU.43c3d.1.P00080146 | Phosphoenolpyruvate carboxykinase, ATP-utilising | scaffold_0008 | 223364 | 225206 | -7.4 |
| PCAU.43c3d.1.P00080164 | Protein kinase-like domain | scaffold_0008 | 260105 | 262500 | -7.0 |
| PCAU.43c3d.1.P00080182 | Sodium/potassium-transporting ATPase signature | scaffold_0008 | 286306 | 290049 | -7.4 |
| PCAU.43c3d.1.P00080185 | TATA-box binding protein | scaffold_0008 | 291411 | 292324 | -6.2 |
| PCAU.43c3d.1.P00080186 | Ribosomal protein L10/acidic P0 | scaffold_0008 | 292437 | 293576 | -7.2 |
| PCAU.43c3d.1.P00080196 | Sodium/potassium-transporting ATPase signature | scaffold_0008 | 304921 | 308602 | -7.0 |
| PCAU.43c3d.1.P00080245 | Histone deacetylase superfamily | scaffold_0008 | 379335 | 380864 | -6.5 |
| PCAU.43c3d.1.P00080298 | Ribosomal protein L7A/L8 | scaffold_0008 | 462850 | 463769 | -6.3 |
| PCAU.43c3d.1.P00090001 | 60S ribosomal protein L6E | scaffold_0009 | 535 | 1232 | -6.2 |
| PCAU.43c3d.1.P00090003 | Translation protein, beta-barrel domain | scaffold_0009 | 4677 | 5990 | -6.5 |
| PCAU.43c3d.1.P00090009 | ATPase, V1 complex, subunit F, eukaryotic | scaffold_0009 | 15172 | 15670 | -5.8 |
| PCAU.43c3d.1.P00090023 | P-loop containing nucleoside triphosphate hydrolase | scaffold_0009 | 30899 | 31588 | -6.4 |
| PCAU.43c3d.1.P00090068 | Ribosomal protein L22/L17, eukaryotic/archaeal | scaffold_0009 | 106543 | 107206 | -6.2 |
| PCAU.43c3d.1.P00090106 | Cysteine alpha-hairpin motif superfamily | scaffold_0009 | 154310 | 154486 | -5.5 |
| PCAU.43c3d.1.P00090113 | Ribosomal protein L22/L17, eukaryotic/archaeal | scaffold_0009 | 160575 | 161244 | -6.1 |
| PCAU.43c3d.1.P00090115 | EF-hand domain pair | scaffold_0009 | 162482 | 163063 | -6.1 |
| PCAU.43c3d.1.P00090236 | Thiolase-like | scaffold_0009 | 361971 | 363321 | -6.9 |
| PCAU.43c3d.1.P00090242 | Adenosylhomocysteinase | scaffold_0009 | 372323 | 373916 | -6.9 |
| PCAU.43c3d.1.P00090254 | Aromatic amino acid hydroxylase | scaffold_0009 | 387275 | 388686 | -8.6 |
| PCAU.43c3d.1.P00090266 | Nucleophile aminohydrolases, N-terminal | scaffold_0009 | 407369 | 408252 | -6.1 |
| PCAU.43c3d.1.P00090274 | Histone deacetylase superfamily | scaffold_0009 | 416559 | 417959 | -6.2 |
| PCAU.43c3d.1.P00090286 | Ribosomal protein L30, ferredoxin-like fold domain | scaffold_0009 | 442375 | 443253 | -5.9 |
| PCAU.43c3d.1.P00100020 | Ribosomal protein S17e | scaffold_0010 | 26682 | 27186 | -6.3 |
| PCAU.43c3d.1.P00100035 | Glutathione S-transferase, C-terminal-like | scaffold_0010 | 47114 | 47897 | -7.7 |
| PCAU.43c3d.1.P00100037 | P-loop containing nucleoside triphosphate hydrolase | scaffold_0010 | 49517 | 50277 | -5.9 |
| PCAU.43c3d.1.P00100039 | Acyl-CoA dehydrogenase/oxidase C-terminal | scaffold_0010 | 52066 | 54145 | -8.2 |
| PCAU.43c3d.1.P00100094 | Ribosomal protein L18e/L15P | scaffold_0010 | 134191 | 134722 | -5.5 |
| PCAU.43c3d.1.P00100130 | Aspartic peptidase | scaffold_0010 | 187402 | 188608 | -7.1 |
| PCAU.43c3d.1.P00100133 | Ribosomal protein L7A/L8 | scaffold_0010 | 190345 | 191272 | -5.8 |
| PCAU.43c3d.1.P00100150 | Acyl-CoA N-acyltransferase | scaffold_0010 | 214488 | 215073 | -6.6 |
| PCAU.43c3d.1.P00100189 | Glycoside hydrolase, family 13 | scaffold_0010 | 263463 | 265057 | -6.7 |
| PCAU.43c3d.1.P00100209 | Ubiquitin-conjugating enzyme/RWD-like | scaffold_0010 | 300758 | 301393 | -6.5 |
| PCAU.43c3d.1.P00100211 | small GTPase Rab1 family profile. | scaffold_0010 | 303666 | 304247 | -6.7 |
| PCAU.43c3d.1.P00100308 | P-loop containing nucleoside triphosphate hydrolase | scaffold_0010 | 453021 | 453666 | -8.2 |

| | | | | | |
|---|---|---|---|---|---|
| PCAU.43c3d.1.P00100310 | Ribosomal protein S19/S15 | scaffold_0010 | 456428 | 456985 | -5.6 |
| PCAU.43c3d.1.P00110008 | Terpenoid synthase | scaffold_0011 | 6967 | 8092 | -6.7 |
| PCAU.43c3d.1.P00110040 | Nucleophile aminohydrolases, N-terminal | scaffold_0011 | 48702 | 49550 | -6.2 |
| PCAU.43c3d.1.P00110060 | Ureohydrolase domain | scaffold_0011 | 75822 | 76774 | -6.0 |
| PCAU.43c3d.1.P00110063 | P-loop containing nucleoside triphosphate hydrolase | scaffold_0011 | 85990 | 88069 | -5.8 |
| PCAU.43c3d.1.P00110080 | P-loop containing nucleoside triphosphate hydrolase | scaffold_0011 | 107929 | 109350 | -6.6 |
| PCAU.43c3d.1.P00110094 | Ribosomal protein L18e/L15P | scaffold_0011 | 130914 | 131446 | -5.6 |
| PCAU.43c3d.1.P00110134 | Isocitrate and isopropylmalate dehydrogenases family | scaffold_0011 | 185812 | 187015 | -6.6 |
| PCAU.43c3d.1.P00110161 | P-loop containing nucleoside triphosphate hydrolase | scaffold_0011 | 232217 | 232976 | -7.1 |
| PCAU.43c3d.1.P00110177 | Ribosomal protein L29e | scaffold_0011 | 253736 | 254018 | -5.5 |
| PCAU.43c3d.1.P00110250 | Tetratricopeptide repeat-containing domain | scaffold_0011 | 362383 | 363992 | -6.7 |
| PCAU.43c3d.1.P00120020 | Adenylosuccinate synthetase | scaffold_0012 | 33606 | 34954 | -7.0 |
| PCAU.43c3d.1.P00120048 | TatD family | scaffold_0012 | 73774 | 74704 | -6.6 |
| PCAU.43c3d.1.P00120051 | HD domain | scaffold_0012 | 78736 | 79316 | -6.1 |
| PCAU.43c3d.1.P00120101 | NADH:cytochrome b5 reductase (CBR) | scaffold_0012 | 144041 | 144844 | -6.5 |
| PCAU.43c3d.1.P00120109 | Ribosomal protein L32e | scaffold_0012 | 156472 | 156961 | -5.7 |
| PCAU.43c3d.1.P00120115 | Peptidase C26, gamma-glutamyl hydrolase | scaffold_0012 | 166815 | 167854 | -6.9 |
| PCAU.43c3d.1.P00120152 | 2-oxoglutarate dehydrogenase E1 component | scaffold_0012 | 232460 | 235571 | -7.3 |
| PCAU.43c3d.1.P00120158 | Ribosomal protein S10 | scaffold_0012 | 241365 | 241823 | -5.3 |
| PCAU.43c3d.1.P00120179 | RNA polymerase, Rpb8 | scaffold_0012 | 273434 | 273975 | -7.3 |
| PCAU.43c3d.1.P00120211 | Ribosomal protein L21-like | scaffold_0012 | 322340 | 323009 | -5.9 |
| PCAU.43c3d.1.P00120232 | ARP2/3 complex, 21kDa subunit (p21-Arc) | scaffold_0012 | 364059 | 364693 | -6.9 |
| PCAU.43c3d.1.P00120273 | Peptidase M16 domain | scaffold_0012 | 409005 | 411915 | -7.7 |
| PCAU.43c3d.1.P00120281 | Metallo-dependent phosphatase-like | scaffold_0012 | 422916 | 423958 | -6.2 |
| PCAU.43c3d.1.P00130003 | Ribosomal protein S15 | scaffold_0013 | 9358 | 9960 | -6.2 |
| PCAU.43c3d.1.P00130042 | Protein kinase-like domain | scaffold_0013 | 71496 | 72759 | -7.9 |
| PCAU.43c3d.1.P00130061 | Coenzyme A transferase family I | scaffold_0013 | 100328 | 101949 | -7.2 |
| PCAU.43c3d.1.P00130064 | DNA replication licensing factor Mcm6 | scaffold_0013 | 104187 | 106641 | -7.0 |
| PCAU.43c3d.1.P00130082 | Peptidase M28 | scaffold_0013 | 130211 | 131683 | -6.5 |
| PCAU.43c3d.1.P00130092 | Protein kinase domain | scaffold_0013 | 140125 | 141136 | -6.4 |
| PCAU.43c3d.1.P00130094 | Ribosomal protein S8 | scaffold_0013 | 143724 | 144231 | -5.5 |
| PCAU.43c3d.1.P00130099 | Dimeric alpha-beta barrel | scaffold_0013 | 149043 | 150107 | -6.7 |
| PCAU.43c3d.1.P00130123 | Ribosomal protein S12e | scaffold_0013 | 193449 | 193929 | -6.4 |
| PCAU.43c3d.1.P00130192 | Protein kinase-like domain | scaffold_0013 | 303106 | 304542 | -6.6 |
| PCAU.43c3d.1.P00130201 | Ribosomal protein L10e/L16 | scaffold_0013 | 314827 | 315583 | -6.1 |
| PCAU.43c3d.1.P00130213 | NADH:cytochrome b5 reductase (CBR) | scaffold_0013 | 332026 | 332918 | -6.6 |
| PCAU.43c3d.1.P00130263 | Nucleoside diphosphate kinase | scaffold_0013 | 395955 | 396503 | -6.0 |
| PCAU.43c3d.1.P00130265 | Nucleoside diphosphate kinase | scaffold_0013 | 397234 | 397748 | -6.4 |
| PCAU.43c3d.1.P00130276 | Nucleic acid-binding, OB-fold | scaffold_0013 | 416785 | 417366 | -6.1 |
| PCAU.43c3d.1.P00140016 | Peptidase C1A | scaffold_0014 | 28067 | 29103 | -6.3 |
| PCAU.43c3d.1.P00140039 | P-type ATPase, transmembrane domain | scaffold_0014 | 56088 | 59625 | -6.7 |
| PCAU.43c3d.1.P00140048 | Serine/threonine/dual specificity protein kinase, catalytic domain | scaffold_0014 | 70894 | 72107 | -6.8 |
| PCAU.43c3d.1.P00140054 | Isocitrate dehydrogenase NADP-dependent | scaffold_0014 | 77490 | 78885 | -6.9 |
| PCAU.43c3d.1.P00140056 | Protein kinase-like domain | scaffold_0014 | 80595 | 81677 | -7.4 |
| PCAU.43c3d.1.P00140111 | Tryptophan-tRNA ligase | scaffold_0014 | 171404 | 172707 | -7.0 |
| PCAU.43c3d.1.P00140120 | Ham1-like protein | scaffold_0014 | 185199 | 185889 | -5.6 |
| PCAU.43c3d.1.P00140144 | Nucleophile aminohydrolases, N-terminal | scaffold_0014 | 220137 | 220943 | -5.9 |
| PCAU.43c3d.1.P00140148 | Ubiquitin-conjugating enzyme/RWD-like | scaffold_0014 | 224507 | 225228 | -6.6 |
| PCAU.43c3d.1.P00140158 | Electron transfer flavoprotein-ubiquinone oxidoreductase | scaffold_0014 | 233271 | 235065 | -7.3 |
| PCAU.43c3d.1.P00140222 | Translation initiation factor IF- 2 | scaffold_0014 | 313354 | 315353 | -6.8 |
| PCAU.43c3d.1.P00140233 | Peptidase C26, gamma-glutamyl hydrolase | scaffold_0014 | 326416 | 327417 | -6.4 |
| PCAU.43c3d.1.P00140258 | Peptidyl-prolyl cis-trans isomerase, PpiC-type | scaffold_0014 | 375609 | 376044 | -5.3 |
| PCAU.43c3d.1.P00150004 | DNA primase, small subunit, eukaryotic/archaeal | scaffold_0015 | 6477 | 7732 | -7.2 |
| PCAU.43c3d.1.P00150008 | P-loop containing nucleoside triphosphate hydrolase | scaffold_0015 | 13295 | 14052 | -6.0 |
| PCAU.43c3d.1.P00150011 | Phosphoribosyltransferase-like | scaffold_0015 | 17607 | 18520 | -6.8 |
| PCAU.43c3d.1.P00150021 | Translation initiation factor IF2/IF5 | scaffold_0015 | 33247 | 33910 | -6.5 |
| PCAU.43c3d.1.P00150095 | Pseudouridine synthase I, TruA | scaffold_0015 | 123163 | 124256 | -6.0 |
| PCAU.43c3d.1.P00150098 | Glutamine synthetase/guanido kinase, catalytic domain | scaffold_0015 | 125433 | 126645 | -7.0 |
| PCAU.43c3d.1.P00150104 | Protein kinase-like domain | scaffold_0015 | 132724 | 133943 | -8.4 |
| PCAU.43c3d.1.P00150183 | Glycine cleavage system T protein | scaffold_0015 | 247866 | 249159 | -7.2 |
| PCAU.43c3d.1.P00150189 | Thiolase-like, subgroup | scaffold_0015 | 254502 | 255812 | -7.8 |
| PCAU.43c3d.1.P00150190 | ATPase, V1 complex, subunit C | scaffold_0015 | 255873 | 257193 | -6.3 |
| PCAU.43c3d.1.P00150202 | Fructose-bisphosphate aldolase class-I, eukaryotic-type | scaffold_0015 | 269083 | 270315 | -6.5 |
| PCAU.43c3d.1.P00150213 | HAD-like domain | scaffold_0015 | 283771 | 285025 | -6.7 |
| PCAU.43c3d.1.P00150236 | Pyridine nucleotide disulphide reductase class-I signature | scaffold_0015 | 323818 | 325375 | -6.7 |
| PCAU.43c3d.1.P00150250 | small GTPase Rab1 family profile. | scaffold_0015 | 358104 | 358723 | -6.1 |
| PCAU.43c3d.1.P00150255 | Phosphoribosyltransferase-like | scaffold_0015 | 363798 | 364661 | -6.2 |
| PCAU.43c3d.1.P00150260 | Protein kinase-like domain | scaffold_0015 | 368455 | 369651 | -8.1 |
| PCAU.43c3d.1.P00150277 | Zinc finger, TFIIS-type | scaffold_0015 | 392631 | 393016 | -6.7 |
| PCAU.43c3d.1.P00160003 | Catalase-like domain | scaffold_0016 | 2433 | 3997 | -6.5 |
| PCAU.43c3d.1.P00160039 | Pyruvate dehydrogenase E1 component subunit beta | scaffold_0016 | 64592 | 65754 | -6.5 |
| PCAU.43c3d.1.P00160047 | Translation protein SH3-like domain | scaffold_0016 | 74553 | 75088 | -6.7 |
| PCAU.43c3d.1.P00160054 | Deoxynucleoside kinase | scaffold_0016 | 86169 | 86896 | -7.2 |
| PCAU.43c3d.1.P00160056 | Isocitrate lyase | scaffold_0016 | 89205 | 90872 | -6.9 |
| PCAU.43c3d.1.P00160160 | Ribosomal protein S25 | scaffold_0016 | 270834 | 271385 | -6.0 |
| PCAU.43c3d.1.P00160192 | Adenylate kinase/UMP-CMP kinase | scaffold_0016 | 312291 | 313154 | -6.5 |
| PCAU.43c3d.1.P00160209 | Zinc-binding ribosomal protein | scaffold_0016 | 339938 | 340596 | -5.9 |
| PCAU.43c3d.1.P00160216 | Peptidase C15, pyroglutamyl peptidase I-like | scaffold_0016 | 347117 | 347746 | -6.8 |
| PCAU.43c3d.1.P00170031 | Autophagy protein Atg8 ubiquitin like | scaffold_0017 | 33563 | 33998 | -7.4 |
| PCAU.43c3d.1.P00170078 | Metallo-dependent phosphatase-like | scaffold_0017 | 109539 | 111245 | -6.3 |
| PCAU.43c3d.1.P00170080 | Ribosomal protein S28e | scaffold_0017 | 112270 | 112528 | -4.8 |
| PCAU.43c3d.1.P00170095 | Aspartic peptidase | scaffold_0017 | 131567 | 132879 | -6.5 |
| PCAU.43c3d.1.P00170098 | Myristoyl-CoA:protein N-myristoyltransferase | scaffold_0017 | 134761 | 136178 | -7.7 |
| PCAU.43c3d.1.P00170105 | Glycosyl transferase, family 35 | scaffold_0017 | 143219 | 146010 | -7.9 |
| PCAU.43c3d.1.P00170130 | Protein kinase-like domain | scaffold_0017 | 175764 | 176918 | -7.1 |
| PCAU.43c3d.1.P00170133 | Ribosomal protein L26/L24P, eukaryotic/archaeal | scaffold_0017 | 179179 | 179730 | -6.0 |
| PCAU.43c3d.1.P00170138 | Protein kinase-like domain | scaffold_0017 | 187845 | 189304 | -7.2 |
| PCAU.43c3d.1.P00170179 | small GTPase Rab1 family profile. | scaffold_0017 | 246572 | 247244 | -6.3 |

| | | | | | |
|---|---|---|---|---|---|
| PCAU.43c3d.1.P00170205 | Peptidase C12, ubiquitin carboxyl-terminal hydrolase | scaffold_0017 | 291231 | 292011 | -7.4 |
| PCAU.43c3d.1.P00170238 | P-type ATPase, transmembrane domain | scaffold_0017 | 355481 | 358910 | -7.0 |
| PCAU.43c3d.1.P00180020 | Ribosomal protein S4/S9 | scaffold_0018 | 19708 | 20368 | -5.6 |
| PCAU.43c3d.1.P00180078 | GHMP kinase, C-terminal domain | scaffold_0018 | 82561 | 82983 | -5.9 |
| PCAU.43c3d.1.P00180097 | Chaperonin Cpn60/TCP-1 | scaffold_0018 | 104030 | 105835 | -5.6 |
| PCAU.43c3d.1.P00180113 | Ribosomal protein L18a/LX | scaffold_0018 | 121623 | 122372 | -6.2 |
| PCAU.43c3d.1.P00180151 | Protein kinase-like domain | scaffold_0018 | 174789 | 177471 | -6.9 |
| PCAU.43c3d.1.P00180156 | Ribosomal protein L5 domain | scaffold_0018 | 183976 | 184678 | -6.2 |
| PCAU.43c3d.1.P00180172 | DNA mismatch repair protein MutS, core | scaffold_0018 | 213468 | 215895 | -6.9 |
| PCAU.43c3d.1.P00180178 | Protein kinase-like domain | scaffold_0018 | 222844 | 223872 | -7.4 |
| PCAU.43c3d.1.P00180197 | small GTPase Rab1 family profile. | scaffold_0018 | 242201 | 243037 | -7.3 |
| PCAU.43c3d.1.P00180208 | Cytochrome c oxidase copper chaperone | scaffold_0018 | 257635 | 257808 | -5.9 |
| PCAU.43c3d.1.P00180227 | Ubiquitin-conjugating enzyme/RWD-like | scaffold_0018 | 282244 | 282992 | -6.0 |
| PCAU.43c3d.1.P00180233 | Nucleophile aminohydrolases, N-terminal | scaffold_0018 | 289166 | 290165 | -6.1 |
| PCAU.43c3d.1.P00180261 | Sodium/potassium-transporting ATPase signature | scaffold_0018 | 336928 | 340800 | -7.5 |
| PCAU.43c3d.1.P00180267 | P-loop containing nucleoside triphosphate hydrolase | scaffold_0018 | 354333 | 355142 | -6.5 |
| PCAU.43c3d.1.P00190001 | ABC transporter type 1, transmembrane domain | scaffold_0019 | 1501 | 6174 | -7.0 |
| PCAU.43c3d.1.P00190011 | 14-3-3 domain | scaffold_0019 | 26362 | 27282 | -5.9 |
| PCAU.43c3d.1.P00190036 | 14-3-3 domain | scaffold_0019 | 74083 | 74963 | -6.0 |
| PCAU.43c3d.1.P00190058 | Ribosomal protein S17e | scaffold_0019 | 114006 | 114508 | -6.4 |
| PCAU.43c3d.1.P00190082 | Ribosomal protein L18e/L15P | scaffold_0019 | 144281 | 144808 | -5.6 |
| PCAU.43c3d.1.P00190084 | Thiamin pyrophosphokinase, catalytic domain | scaffold_0019 | 145979 | 146647 | -7.1 |
| PCAU.43c3d.1.P00190114 | ATPase, V1/A1 complex, subunit E | scaffold_0019 | 206450 | 207272 | -6.2 |
| PCAU.43c3d.1.P00190125 | P-loop containing nucleoside triphosphate hydrolase | scaffold_0019 | 221591 | 222350 | -6.0 |
| PCAU.43c3d.1.P00190165 | Protein kinase-like domain | scaffold_0019 | 292595 | 293868 | -6.8 |
| PCAU.43c3d.1.P00190189 | Ribosomal protein L18e | scaffold_0019 | 327742 | 328418 | -7.1 |
| PCAU.43c3d.1.P00200036 | Sugar-phosphate isomerase, RpiB/LacA/LacB family | scaffold_0020 | 53982 | 54512 | -5.5 |
| PCAU.43c3d.1.P00200037 | Autophagy protein Atg8 ubiquitin like | scaffold_0020 | 54497 | 54911 | -5.3 |
| PCAU.43c3d.1.P00200043 | Cyclic nucleotide-binding domain | scaffold_0020 | 64410 | 66999 | -7.5 |
| PCAU.43c3d.1.P00200057 | Aminotransferase, class IV | scaffold_0020 | 82453 | 83880 | -6.7 |
| PCAU.43c3d.1.P00200078 | Peptidase C13, legumain | scaffold_0020 | 109401 | 110751 | -7.1 |
| PCAU.43c3d.1.P00200090 | P-loop containing nucleoside triphosphate hydrolase | scaffold_0020 | 125916 | 126765 | -6.3 |
| PCAU.43c3d.1.P00200122 | P-loop containing nucleoside triphosphate hydrolase | scaffold_0020 | 173564 | 174646 | -7.4 |
| PCAU.43c3d.1.P00200147 | Translation Initiation factor eIF- 4e-like domain | scaffold_0020 | 217759 | 218366 | -5.6 |
| PCAU.43c3d.1.P00200158 | Ribosomal protein L13e | scaffold_0020 | 235281 | 236015 | -6.0 |
| PCAU.43c3d.1.P00200159 | Cyclic nucleotide-binding domain | scaffold_0020 | 236005 | 238574 | -7.2 |
| PCAU.43c3d.1.P00200181 | Nucleophile aminohydrolases, N-terminal | scaffold_0020 | 273016 | 273699 | -5.8 |
| PCAU.43c3d.1.P00200193 | Ran GTPase | scaffold_0020 | 289398 | 290160 | -7.1 |
| PCAU.43c3d.1.P00200196 | Cytidyltransferase-like domain | scaffold_0020 | 293410 | 294046 | -5.6 |
| PCAU.43c3d.1.P00200197 | Ribosomal protein L36e | scaffold_0020 | 293997 | 294383 | -5.6 |
| PCAU.43c3d.1.P00210012 | Protein translocase SecE domain | scaffold_0021 | 16142 | 16390 | -5.3 |
| PCAU.43c3d.1.P00210015 | P-loop containing nucleoside triphosphate hydrolase | scaffold_0021 | 21028 | 22319 | -6.2 |
| PCAU.43c3d.1.P00210017 | small GTPase Rab1 family profile. | scaffold_0021 | 22713 | 23480 | -6.8 |
| PCAU.43c3d.1.P00210034 | Glutamyl/glutaminyl-tRNA synthetase | scaffold_0021 | 51146 | 53576 | -8.4 |
| PCAU.43c3d.1.P00210076 | P-loop containing nucleoside triphosphate hydrolase | scaffold_0021 | 97287 | 98062 | -6.4 |
| PCAU.43c3d.1.P00210085 | Protein kinase-like domain | scaffold_0021 | 108897 | 109904 | -7.0 |
| PCAU.43c3d.1.P00210094 | Triosephosphate isomerase | scaffold_0021 | 117253 | 118128 | -6.7 |
| PCAU.43c3d.1.P00210095 | small GTPase Rab1 family profile. | scaffold_0021 | 118060 | 118703 | -6.8 |
| PCAU.43c3d.1.P00210115 | Manganese/iron superoxide dismutase | scaffold_0021 | 144534 | 145283 | -6.4 |
| PCAU.43c3d.1.P00210119 | Phospholipase/carboxylesterase/thioesterase | scaffold_0021 | 149392 | 150157 | -6.1 |
| PCAU.43c3d.1.P00210141 | Tetrapyrrole methylase | scaffold_0021 | 183320 | 184212 | -8.3 |
| PCAU.43c3d.1.P00210156 | NUDIX hydrolase domain | scaffold_0021 | 206360 | 206864 | -6.3 |
| PCAU.43c3d.1.P00210176 | small GTPase Rab1 family profile. | scaffold_0021 | 234161 | 234847 | -6.4 |
| PCAU.43c3d.1.P00210186 | P-loop containing nucleoside triphosphate hydrolase | scaffold_0021 | 247225 | 249002 | -7.3 |
| PCAU.43c3d.1.P00210229 | Ribokinase-like | scaffold_0021 | 306035 | 307219 | -6.8 |
| PCAU.43c3d.1.P00220015 | 14-3-3 protein | scaffold_0022 | 20353 | 21120 | -6.4 |
| PCAU.43c3d.1.P00220028 | Fructose-1,6-bisphosphatase class 1/Sedoheputulose-1,7-bisphosphatase | scaffold_0022 | 40370 | 41406 | -6.1 |
| PCAU.43c3d.1.P00220032 | Peptidase M1, alanine aminopeptidase/leukotriene A4 hydrolase | scaffold_0022 | 46163 | 48195 | -8.0 |
| PCAU.43c3d.1.P00220067 | Chaperonin Cpn60/TCP-1 | scaffold_0022 | 109399 | 111186 | -6.4 |
| PCAU.43c3d.1.P00220075 | Ribosomal protein L13e | scaffold_0022 | 120087 | 120837 | -6.0 |
| PCAU.43c3d.1.P00220104 | Electron transfer flavoprotein, beta subunit | scaffold_0022 | 172004 | 172827 | -6.9 |
| PCAU.43c3d.1.P00220135 | Chaperonin Cpn60/TCP-1 | scaffold_0022 | 224198 | 226020 | -5.6 |
| PCAU.43c3d.1.P00220137 | Succinyl-CoA synthetase, beta subunit | scaffold_0022 | 228148 | 229536 | -6.4 |
| PCAU.43c3d.1.P00220155 | Ornithine decarboxylase | scaffold_0022 | 253105 | 254561 | -7.2 |
| PCAU.43c3d.1.P00220229 | Protein kinase-like domain | scaffold_0022 | 358410 | 359518 | -7.1 |
| PCAU.43c3d.1.P00230005 | JAB1/MPN/MOV34 metalloenzyme domain | scaffold_0023 | 10470 | 11489 | -6.3 |
| PCAU.43c3d.1.P00230014 | P-loop containing nucleoside triphosphate hydrolase | scaffold_0023 | 24562 | 25239 | -6.2 |
| PCAU.43c3d.1.P00230061 | Ran GTPase | scaffold_0023 | 142052 | 142852 | -7.3 |
| PCAU.43c3d.1.P00230062 | JAB1/MPN/MOV34 metalloenzyme domain | scaffold_0023 | 142946 | 144009 | -6.8 |
| PCAU.43c3d.1.P00230063 | Serine/threonine/dual specificity protein kinase, catalytic domain | scaffold_0023 | 144034 | 144976 | -7.0 |
| PCAU.43c3d.1.P00230065 | Ribosomal protein L38e | scaffold_0023 | 146723 | 147119 | -6.2 |
| PCAU.43c3d.1.P00230066 | Casein kinase II, regulatory subunit | scaffold_0023 | 147115 | 147950 | -6.4 |
| PCAU.43c3d.1.P00230077 | Acyl-CoA dehydrogenase/oxidase C-terminal | scaffold_0023 | 160722 | 162620 | -7.8 |
| PCAU.43c3d.1.P00230080 | DHS-like NAD/FAD-binding domain | scaffold_0023 | 164257 | 165323 | -5.9 |
| PCAU.43c3d.1.P00230098 | Initiation factor 2B-related | scaffold_0023 | 192189 | 193613 | -7.5 |
| PCAU.43c3d.1.P00230108 | small GTPase Rab1 family profile. | scaffold_0023 | 216541 | 217261 | -6.3 |
| PCAU.43c3d.1.P00230123 | Ribonuclease H-like domain | scaffold_0023 | 232132 | 234559 | -7.5 |
| PCAU.43c3d.1.P00230127 | V-type ATPase, V0 complex, 116kDa subunit family | scaffold_0023 | 241389 | 243879 | -7.1 |
| PCAU.43c3d.1.P00230130 | Ribosomal protein L5 | scaffold_0023 | 244920 | 245634 | -6.1 |
| PCAU.43c3d.1.P00230131 | Ribosomal protein S3Ae | scaffold_0023 | 245733 | 246624 | -5.5 |
| PCAU.43c3d.1.P00230168 | Peptidase M1, alanine aminopeptidase/leukotriene A4 hydrolase | scaffold_0023 | 297131 | 299171 | -7.5 |
| PCAU.43c3d.1.P00240039 | Thioredoxin domain | scaffold_0024 | 62490 | 64073 | -7.4 |
| PCAU.43c3d.1.P00240046 | Thiolase-like, subgroup | scaffold_0024 | 75494 | 76816 | -6.3 |
| PCAU.43c3d.1.P00240061 | Translation release factor pelota | scaffold_0024 | 103079 | 104449 | -7.0 |
| PCAU.43c3d.1.P00240064 | Ribosomal protein L10/acidic P0 | scaffold_0024 | 107238 | 108343 | -5.7 |
| PCAU.43c3d.1.P00240084 | Ribosomal protein S19e | scaffold_0024 | 135660 | 136219 | -5.7 |
| PCAU.43c3d.1.P00240093 | Citrate synthase-like | scaffold_0024 | 146994 | 148243 | -6.2 |
| PCAU.43c3d.1.P00240104 | Thiamin diphosphate-binding fold | scaffold_0024 | 166976 | 168347 | -6.7 |

| | | | | | |
|---|---|---|---|---|---|
| PCAU.43c3d.1.P00240168 | Phosphoenolpyruvate carboxykinase, ATP-utilising | scaffold_0024 | 261971 | 263799 | -7.2 |
| PCAU.43c3d.1.P00240171 | Ribosomal protein L29 | scaffold_0024 | 266099 | 266583 | -5.8 |
| PCAU.43c3d.1.P00240178 | Heat shock protein Hsp90 family | scaffold_0024 | 276101 | 278218 | -6.1 |
| PCAU.43c3d.1.P00240180 | Ribosomal protein S7e | scaffold_0024 | 279276 | 280025 | -6.1 |
| PCAU.43c3d.1.P00240230 | Ribosomal protein L35A | scaffold_0024 | 343013 | 343510 | -5.6 |
| PCAU.43c3d.1.P00250008 | Ribosomal protein S4/S9 | scaffold_0025 | 9106 | 9770 | -5.8 |
| PCAU.43c3d.1.P00250044 | small GTPase Rab1 family profile. | scaffold_0025 | 58656 | 59390 | -6.5 |
| PCAU.43c3d.1.P00250065 | Tyrosine-tRNA ligase, bacterial-type | scaffold_0025 | 80649 | 82233 | -6.8 |
| PCAU.43c3d.1.P00250081 | Protein kinase-like domain | scaffold_0025 | 105270 | 106159 | -6.3 |
| PCAU.43c3d.1.P00250203 | Metal-dependent hydrolase, composite domain | scaffold_0025 | 311952 | 313256 | -6.2 |
| PCAU.43c3d.1.P00250221 | Ribosomal protein L5 eukaryotic/L18 archaeal | scaffold_0025 | 336092 | 337122 | -6.1 |
| PCAU.43c3d.1.P00250227 | Tetratricopeptide-like helical domain | scaffold_0025 | 343134 | 344669 | -6.9 |
| PCAU.43c3d.1.P00260014 | Ribosomal protein S28e | scaffold_0026 | 21077 | 21356 | -4.8 |
| PCAU.43c3d.1.P00260017 | Protein kinase-like domain | scaffold_0026 | 23309 | 24374 | -6.6 |
| PCAU.43c3d.1.P00260061 | Cyclic nucleotide-binding-like | scaffold_0026 | 83286 | 85769 | -7.6 |
| PCAU.43c3d.1.P00260071 | Ribosomal protein L26/L24P, eukaryotic/archaeal | scaffold_0026 | 101704 | 102229 | -6.1 |
| PCAU.43c3d.1.P00260122 | Cyclic nucleotide-binding domain | scaffold_0026 | 172826 | 174002 | -6.0 |
| PCAU.43c3d.1.P00260129 | P-loop containing nucleoside triphosphate hydrolase | scaffold_0026 | 178574 | 179386 | -8.0 |
| PCAU.43c3d.1.P00260145 | Phosphoglycerate mutase 1 | scaffold_0026 | 204662 | 205511 | -6.8 |
| PCAU.43c3d.1.P00260148 | Protein kinase domain | scaffold_0026 | 207798 | 208807 | -6.4 |
| PCAU.43c3d.1.P00260169 | Aspartic peptidase | scaffold_0026 | 239458 | 240809 | -6.6 |
| PCAU.43c3d.1.P00260220 | small GTPase Rab1 family profile. | scaffold_0026 | 317487 | 318301 | -6.4 |
| PCAU.43c3d.1.P00260229 | Thioredoxin-like fold | scaffold_0026 | 329973 | 330791 | -6.2 |
| PCAU.43c3d.1.P00260232 | Histidine-tRNA ligase/ATP phosphoribosyltransferase regulatory subunit | scaffold_0026 | 335765 | 337345 | -7.4 |
| PCAU.43c3d.1.P00270007 | Protein kinase domain | scaffold_0027 | 20910 | 23399 | -6.7 |
| PCAU.43c3d.1.P00270035 | Aminoacyl-tRNA synthetase, class Ic | scaffold_0027 | 59970 | 61284 | -7.1 |
| PCAU.43c3d.1.P00270046 | Peptidase M24, structural domain | scaffold_0027 | 76292 | 77460 | -6.1 |
| PCAU.43c3d.1.P00270065 | Heat shock protein 70 family | scaffold_0027 | 117903 | 119893 | -6.0 |
| PCAU.43c3d.1.P00270098 | Mitochondrial carrier domain | scaffold_0027 | 166822 | 167896 | -7.0 |
| PCAU.43c3d.1.P00270100 | Thiolase-like | scaffold_0027 | 169816 | 171093 | -6.9 |
| PCAU.43c3d.1.P00270153 | Protein kinase-like domain | scaffold_0027 | 266987 | 268271 | -7.4 |
| PCAU.43c3d.1.P00280039 | Chaperonin Cpn10 | scaffold_0028 | 83016 | 83381 | -5.7 |
| PCAU.43c3d.1.P00280097 | Sodium:neurotransmitter symporter | scaffold_0028 | 152086 | 153825 | -7.9 |
| PCAU.43c3d.1.P00280108 | Peptidase M1, alanine aminopeptidase/leukotriene A4 hydrolase | scaffold_0028 | 170592 | 172566 | -7.0 |
| PCAU.43c3d.1.P00280118 | WD40 repeat | scaffold_0028 | 183248 | 184426 | -7.2 |
| PCAU.43c3d.1.P00280162 | Heat shock protein 70 family | scaffold_0028 | 254433 | 256501 | -6.3 |
| PCAU.43c3d.1.P00280167 | Small GTPase superfamily, ARF type | scaffold_0028 | 261742 | 262339 | -6.1 |
| PCAU.43c3d.1.P00280176 | Nucleoside phosphatase GDA1/CD39 | scaffold_0028 | 269735 | 271047 | -7.2 |
| PCAU.43c3d.1.P00280180 | P-loop containing nucleoside triphosphate hydrolase | scaffold_0028 | 276224 | 276940 | -6.4 |
| PCAU.43c3d.1.P00280205 | Anaphase-promoting complex subunit 11 | scaffold_0028 | 328740 | 328985 | -5.8 |
| PCAU.43c3d.1.P00290007 | Ribosome biogenesis factor NIP7-like | scaffold_0029 | 13098 | 13746 | -6.3 |
| PCAU.43c3d.1.P00290015 | Serine/threonine-protein kinase Bud32 | scaffold_0029 | 26836 | 27500 | -6.7 |
| PCAU.43c3d.1.P00290026 | Ribosomal protein L2 | scaffold_0029 | 51972 | 52854 | -7.2 |
| PCAU.43c3d.1.P00290029 | Ribosomal protein L22e | scaffold_0029 | 56907 | 57371 | -6.3 |
| PCAU.43c3d.1.P00290032 | Tubulin | scaffold_0029 | 59712 | 61079 | -6.0 |
| PCAU.43c3d.1.P00290036 | Cyclic nucleotide-binding domain | scaffold_0029 | 65608 | 66816 | -6.2 |
| PCAU.43c3d.1.P00290044 | Ribosomal protein L34Ae | scaffold_0029 | 74427 | 74892 | -5.5 |
| PCAU.43c3d.1.P00290051 | Protein kinase-like domain | scaffold_0029 | 82021 | 82975 | -6.7 |
| PCAU.43c3d.1.P00290073 | S-adenosylmethionine synthetase | scaffold_0029 | 120236 | 121606 | -6.8 |
| PCAU.43c3d.1.P00290090 | RuvB-like | scaffold_0029 | 153336 | 154837 | -6.6 |
| PCAU.43c3d.1.P00290163 | DNA mismatch repair protein MutS-homologue MSH6 | scaffold_0029 | 251459 | 254717 | -7.2 |
| PCAU.43c3d.1.P00290164 | O-phosphoseryl-tRNA(Sec) selenium transferase | scaffold_0029 | 254739 | 256218 | -6.4 |
| PCAU.43c3d.1.P00290178 | Glutamyl/glutaminyl-tRNA synthetase | scaffold_0029 | 273093 | 274956 | -7.0 |
| PCAU.43c3d.1.P00300001 | Ribosomal protein L14b/L23e | scaffold_0030 | 7705 | 8241 | -6.3 |
| PCAU.43c3d.1.P00300023 | Ribosomal protein S10 | scaffold_0030 | 42638 | 43121 | -6.2 |
| PCAU.43c3d.1.P00300050 | P-loop containing nucleoside triphosphate hydrolase | scaffold_0030 | 90550 | 91245 | -6.0 |
| PCAU.43c3d.1.P00300064 | DAHP synthetase, class II | scaffold_0030 | 108522 | 109772 | -6.4 |
| PCAU.43c3d.1.P00300077 | SKP1 component | scaffold_0030 | 123317 | 123934 | -5.8 |
| PCAU.43c3d.1.P00300095 | RIO kinase | scaffold_0030 | 141308 | 142684 | -3.8 |
| PCAU.43c3d.1.P00300102 | L-Aspartase-like | scaffold_0030 | 150592 | 152223 | -6.7 |
| PCAU.43c3d.1.P00300125 | Armadillo-type fold | scaffold_0030 | 182273 | 185188 | -8.4 |
| PCAU.43c3d.1.P00300132 | DNA-directed RNA pol I, largest subunit | scaffold_0030 | 194361 | 199341 | -8.0 |
| PCAU.43c3d.1.P00300137 | Heat shock protein 70 family | scaffold_0030 | 206585 | 209179 | -7.5 |
| PCAU.43c3d.1.P00300149 | Protein kinase-like domain | scaffold_0030 | 235164 | 236558 | -8.7 |
| PCAU.43c3d.1.P00300156 | P-loop containing nucleoside triphosphate hydrolase | scaffold_0030 | 246067 | 247695 | -6.5 |
| PCAU.43c3d.1.P00300190 | Protein kinase-like domain | scaffold_0030 | 291183 | 292243 | -7.0 |
| PCAU.43c3d.1.P00310021 | Ammonium transporter | scaffold_0031 | 27846 | 29370 | -7.1 |
| PCAU.43c3d.1.P00310038 | Nucleophile aminohydrolases, N-terminal | scaffold_0031 | 52875 | 53730 | -6.1 |
| PCAU.43c3d.1.P00310048 | P-loop containing nucleoside triphosphate hydrolase | scaffold_0031 | 63168 | 63984 | -7.2 |
| PCAU.43c3d.1.P00310066 | small GTPase Rab1 family profile. | scaffold_0031 | 90980 | 91757 | -6.5 |
| PCAU.43c3d.1.P00310081 | Ribosomal protein L18a/LX | scaffold_0031 | 109805 | 110554 | -6.8 |
| PCAU.43c3d.1.P00310094 | P-loop containing nucleoside triphosphate hydrolase | scaffold_0031 | 130421 | 130873 | -6.7 |
| PCAU.43c3d.1.P00310119 | Deoxyribonuclease II | scaffold_0031 | 180509 | 181695 | -6.2 |
| PCAU.43c3d.1.P00310160 | Pyridine nucleotide disulphide reductase class-I signature | scaffold_0031 | 246607 | 248713 | -6.4 |
| PCAU.43c3d.1.P00310170 | Cyclin-dependent kinase, regulatory subunit | scaffold_0031 | 260483 | 260782 | -7.3 |
| PCAU.43c3d.1.P00310178 | Ribosomal protein S12/S23 | scaffold_0031 | 266727 | 267245 | -5.8 |
| PCAU.43c3d.1.P00320013 | WD40/YVTN repeat-like-containing domain | scaffold_0032 | 21607 | 23225 | -7.0 |
| PCAU.43c3d.1.P00320017 | Isocitrate dehydrogenase NADP-dependent | scaffold_0032 | 24916 | 26255 | -7.3 |
| PCAU.43c3d.1.P00320108 | Peptidase C1A | scaffold_0032 | 180009 | 180991 | -6.4 |
| PCAU.43c3d.1.P00320114 | P-loop containing nucleoside triphosphate hydrolase | scaffold_0032 | 189079 | 189863 | -5.8 |
| PCAU.43c3d.1.P00320121 | Peptidase C12, ubiquitin carboxyl-terminal hydrolase | scaffold_0032 | 201457 | 202679 | -6.8 |
| PCAU.43c3d.1.P00320142 | Peptidase C1A | scaffold_0032 | 255628 | 256678 | -6.8 |
| PCAU.43c3d.1.P00330006 | MEMO1 family | scaffold_0033 | 7127 | 8200 | -6.3 |
| PCAU.43c3d.1.P00330043 | Mitochondrial carrier domain | scaffold_0033 | 66058 | 67025 | -7.2 |
| PCAU.43c3d.1.P00330049 | Malate dehydrogenase, type 2 | scaffold_0033 | 71418 | 72578 | -7.1 |
| PCAU.43c3d.1.P00330073 | Armadillo-like helical | scaffold_0033 | 97105 | 98024 | -6.8 |
| PCAU.43c3d.1.P00330096 | Metallo-dependent phosphatase-like | scaffold_0033 | 136640 | 138422 | -7.0 |
| PCAU.43c3d.1.P00330101 | Chorismate synthase | scaffold_0033 | 142955 | 144025 | -6.0 |

| | | | | | |
|---|---|---|---|---|---|
| PCAU.43c3d.1.P00330103 | Glycosyl transferase, family 35 | scaffold_0033 | 147181 | 149963 | -7.4 |
| PCAU.43c3d.1.P00330117 | Acylphosphatase-like domain | scaffold_0033 | 168248 | 168655 | -5.4 |
| PCAU.43c3d.1.P00330148 | D-Tyr tRNAtyr deacylase-like domain | scaffold_0033 | 225145 | 225655 | -7.3 |
| PCAU.43c3d.1.P00330166 | Autophagy protein Atg8 ubiquitin like | scaffold_0033 | 272144 | 272604 | -6.1 |
| PCAU.43c3d.1.P00340021 | P-type ATPase, transmembrane domain | scaffold_0034 | 26460 | 29662 | -8.2 |
| PCAU.43c3d.1.P00340030 | Malate dehydrogenase, type 2 | scaffold_0034 | 43394 | 44491 | -7.0 |
| PCAU.43c3d.1.P00340118 | DNA binding domain, putative | scaffold_0034 | 167295 | 169148 | -6.6 |
| PCAU.43c3d.1.P00340148 | Protein kinase-like domain | scaffold_0034 | 209094 | 210280 | -6.3 |
| PCAU.43c3d.1.P00340166 | Peptide methionine sulfoxide reductase | scaffold_0034 | 227506 | 228093 | -5.9 |
| PCAU.43c3d.1.P00340167 | Glycoside hydrolase, superfamily | scaffold_0034 | 228036 | 229293 | -6.5 |
| PCAU.43c3d.1.P00350014 | Uncharacterised protein family UPF0001 | scaffold_0035 | 14704 | 15408 | -6.6 |
| PCAU.43c3d.1.P00350088 | Rossmann-like alpha/beta/alpha sandwich fold | scaffold_0035 | 110081 | 111998 | -7.8 |
| PCAU.43c3d.1.P00350094 | P-loop containing nucleoside triphosphate hydrolase | scaffold_0035 | 122271 | 123097 | -6.9 |
| PCAU.43c3d.1.P00350125 | Protein kinase-like domain | scaffold_0035 | 166226 | 167331 | -6.9 |
| PCAU.43c3d.1.P00350141 | Ribosomal protein S30 | scaffold_0035 | 188138 | 188489 | -7.2 |
| PCAU.43c3d.1.P00350149 | Rossmann-like alpha/beta/alpha sandwich fold | scaffold_0035 | 200821 | 202730 | -7.2 |
| PCAU.43c3d.1.P00350154 | Mevalonate kinase | scaffold_0035 | 208153 | 209027 | -6.6 |
| PCAU.43c3d.1.P00350159 | Ribosomal protein S12/S23 | scaffold_0035 | 213637 | 214177 | -5.8 |
| PCAU.43c3d.1.P00350177 | Protein arginine N-methyltransferase PRMT5 | scaffold_0035 | 246451 | 248320 | -7.4 |
| PCAU.43c3d.1.P00350179 | Nucleophile aminohydrolases, N-terminal | scaffold_0035 | 248638 | 249453 | -7.1 |
| PCAU.43c3d.1.P00350219 | Chaperonin Cpn60/TCP-1 | scaffold_0035 | 293119 | 294874 | -6.3 |
| PCAU.43c3d.1.P00350222 | NADH-quinone oxidoreductase subunit E-like | scaffold_0035 | 296870 | 297757 | -6.4 |
| PCAU.43c3d.1.P00350235 | Adenylate kinase/UMP-CMP kinase | scaffold_0035 | 313053 | 313902 | -7.3 |
| PCAU.43c3d.1.P00350236 | Citrate synthase-like | scaffold_0035 | 314634 | 316007 | -6.5 |
| PCAU.43c3d.1.P00360009 | Thioredoxin-like fold | scaffold_0036 | 21349 | 21890 | -6.1 |
| PCAU.43c3d.1.P00360034 | 14-3-3 domain | scaffold_0036 | 54747 | 55540 | -6.7 |
| PCAU.43c3d.1.P00360052 | Ribosomal protein S25 | scaffold_0036 | 80847 | 81383 | -5.7 |
| PCAU.43c3d.1.P00360094 | small GTPase Rab1 family profile. | scaffold_0036 | 135621 | 136356 | -6.4 |
| PCAU.43c3d.1.P00360119 | P-loop containing nucleoside triphosphate hydrolase | scaffold_0036 | 165577 | 166182 | -6.9 |
| PCAU.43c3d.1.P00360138 | Glycoside hydrolase, family 13 | scaffold_0036 | 189149 | 191454 | -6.8 |
| PCAU.43c3d.1.P00360162 | NADH:ubiquinone oxidoreductase, 51kDa subunit | scaffold_0036 | 220408 | 221958 | -7.2 |
| PCAU.43c3d.1.P00360178 | Tubulin | scaffold_0036 | 246266 | 247576 | -6.6 |
| PCAU.43c3d.1.P00360211 | Ribosomal L28e/Mak16 | scaffold_0036 | 303380 | 303917 | -7.0 |
| PCAU.43c3d.1.P00370023 | Ribosomal protein L37e | scaffold_0037 | 29492 | 29895 | -5.9 |
| PCAU.43c3d.1.P00370039 | Protein kinase-like domain | scaffold_0037 | 55179 | 56479 | -6.7 |
| PCAU.43c3d.1.P00370082 | DNA replication licensing factor Mcm7 | scaffold_0037 | 118505 | 120869 | -8.2 |
| PCAU.43c3d.1.P00370095 | Ubiquitin-related domain | scaffold_0037 | 135938 | 136145 | -6.8 |
| PCAU.43c3d.1.P00370133 | Ribosomal protein L24e-related | scaffold_0037 | 190767 | 191367 | -5.7 |
| PCAU.43c3d.1.P00370142 | Ribosomal protein L11/L12 | scaffold_0037 | 202463 | 203070 | -6.1 |
| PCAU.43c3d.1.P00370143 | DNA/pantothenate metabolism flavoprotein, C-terminal | scaffold_0037 | 203063 | 203924 | -7.8 |
| PCAU.43c3d.1.P00370166 | Peptidase M24, structural domain | scaffold_0037 | 268232 | 270131 | -7.5 |
| PCAU.43c3d.1.P00370183 | Initiation factor 2B-related | scaffold_0037 | 293983 | 295060 | -6.4 |
| PCAU.43c3d.1.P00380040 | Armadillo-type fold | scaffold_0038 | 58871 | 60720 | -5.8 |
| PCAU.43c3d.1.P00380074 | P-loop containing nucleoside triphosphate hydrolase | scaffold_0038 | 116042 | 117353 | -6.1 |
| PCAU.43c3d.1.P00380084 | Glycine cleavage system P protein | scaffold_0038 | 128908 | 131912 | -6.2 |
| PCAU.43c3d.1.P00380087 | Pyridoxal phosphate-dependent transferase | scaffold_0038 | 135133 | 136703 | -6.9 |
| PCAU.43c3d.1.P00380102 | Aldehyde/histidinol dehydrogenase | scaffold_0038 | 158645 | 160244 | -7.6 |
| PCAU.43c3d.1.P00380122 | DNA-directed RNA polymerase, subunit 2 | scaffold_0038 | 200243 | 204000 | -6.4 |
| PCAU.43c3d.1.P00380130 | Ribonuclease H-like domain | scaffold_0038 | 226702 | 229128 | -7.8 |
| PCAU.43c3d.1.P00390011 | Peptidyl-tRNA hydrolase | scaffold_0039 | 18396 | 19034 | -7.3 |
| PCAU.43c3d.1.P00390021 | Protein kinase-like domain | scaffold_0039 | 33204 | 34352 | -7.0 |
| PCAU.43c3d.1.P00390025 | Peptidase M1, alanine aminopeptidase/leukotriene A4 hydrolase | scaffold_0039 | 38393 | 40398 | -7.9 |
| PCAU.43c3d.1.P00390040 | Ubiquitin-related domain | scaffold_0039 | 62843 | 63314 | -5.8 |
| PCAU.43c3d.1.P00390078 | Cyclic nucleotide-binding-like | scaffold_0039 | 117105 | 119707 | -7.7 |
| PCAU.43c3d.1.P00390087 | Ergosterol biosynthesis ERG4/ERG24 | scaffold_0039 | 132871 | 134052 | -6.5 |
| PCAU.43c3d.1.P00390154 | 4-hydroxyphenylpyruvate dioxygenase | scaffold_0039 | 229561 | 230857 | -6.3 |
| PCAU.43c3d.1.P00390207 | DNA-directed RNA polymerase III subunit RPC1 | scaffold_0039 | 299630 | 303805 | -6.8 |
| PCAU.43c3d.1.P00400024 | Cyclic nucleotide-binding domain | scaffold_0040 | 34433 | 36839 | -7.5 |
| PCAU.43c3d.1.P00400106 | Tensin phosphatase, lipid phosphatase domain | scaffold_0040 | 184512 | 185569 | -6.3 |
| PCAU.43c3d.1.P00400151 | P-loop containing nucleoside triphosphate hydrolase | scaffold_0040 | 261266 | 262587 | -7.4 |
| PCAU.43c3d.1.P00400175 | Methylpurine-DNA glycosylase (MPG) | scaffold_0040 | 297150 | 298064 | -6.6 |
| PCAU.43c3d.1.P00410005 | Heat shock protein 70 family | scaffold_0041 | 6372 | 8525 | -6.6 |
| PCAU.43c3d.1.P00410013 | AMP-dependent synthetase/ligase | scaffold_0041 | 18843 | 20883 | -6.4 |
| PCAU.43c3d.1.P00410037 | ClpP/crotonase-like domain | scaffold_0041 | 54126 | 55300 | -6.6 |
| PCAU.43c3d.1.P00410050 | Ribosomal protein S12e | scaffold_0041 | 70058 | 70540 | -5.6 |
| PCAU.43c3d.1.P00410058 | DNA-directed RNA polymerase, subunit 2 | scaffold_0041 | 79598 | 80491 | -7.5 |
| PCAU.43c3d.1.P00410074 | Histidine phosphatase superfamily | scaffold_0041 | 105098 | 106494 | -6.9 |
| PCAU.43c3d.1.P00410104 | Armadillo-like helical | scaffold_0041 | 150559 | 152467 | -6.1 |
| PCAU.43c3d.1.P00410112 | NAD(P)-binding domain | scaffold_0041 | 159226 | 160096 | -7.3 |
| PCAU.43c3d.1.P00410118 | Succinyl-CoA synthetase, beta subunit | scaffold_0041 | 164397 | 165807 | -6.5 |
| PCAU.43c3d.1.P00410153 | Peptidase M18 | scaffold_0041 | 221194 | 222696 | -6.2 |
| PCAU.43c3d.1.P00410181 | Thioredoxin-like fold | scaffold_0041 | 264238 | 264983 | -5.8 |
| PCAU.43c3d.1.P00410195 | Protein phosphatase 2A, regulatory subunit PR55 | scaffold_0041 | 283138 | 284505 | -7.0 |
| PCAU.43c3d.1.P00410197 | Ribosomal protein S27e | scaffold_0041 | 285550 | 285922 | -5.9 |
| PCAU.43c3d.1.P00420027 | Protein kinase-like domain | scaffold_0042 | 37233 | 38303 | -6.5 |
| PCAU.43c3d.1.P00420028 | Zinc finger, RING/FYVE/PHD-type | scaffold_0042 | 38289 | 38630 | -5.9 |
| PCAU.43c3d.1.P00420029 | Protein kinase-like domain | scaffold_0042 | 38641 | 39718 | -7.2 |
| PCAU.43c3d.1.P00420056 | Ribosomal protein L27e | scaffold_0042 | 67739 | 68341 | -5.8 |
| PCAU.43c3d.1.P00420112 | Peptidyl-prolyl cis-trans isomerase, FKBP-type | scaffold_0042 | 157638 | 158037 | -6.8 |
| PCAU.43c3d.1.P00420113 | TATA-box binding protein | scaffold_0042 | 158071 | 158731 | -5.8 |
| PCAU.43c3d.1.P00420132 | Thioredoxin-like fold | scaffold_0042 | 196722 | 198348 | -7.1 |
| PCAU.43c3d.1.P00420136 | Enolase | scaffold_0042 | 201298 | 202756 | -6.3 |
| PCAU.43c3d.1.P00420167 | EF-hand domain pair | scaffold_0042 | 248271 | 248826 | -6.2 |
| PCAU.43c3d.1.P00420191 | Peptidase M24, structural domain | scaffold_0042 | 284009 | 285100 | -7.2 |
| PCAU.43c3d.1.P00420195 | Peptidase M16 domain | scaffold_0042 | 287991 | 289456 | -7.2 |
| PCAU.43c3d.1.P00420197 | EF-hand domain | scaffold_0042 | 291199 | 291735 | -8.2 |
| PCAU.43c3d.1.P00430023 | Metallo-dependent phosphatase-like | scaffold_0043 | 26603 | 27689 | -6.6 |
| PCAU.43c3d.1.P00430025 | Peptidase C65, otubain | scaffold_0043 | 27941 | 28843 | -6.2 |

| | | | | | |
|---|---|---|---|---|---|
| PCAU.43c3d.1.P00430028 | Cyclic nucleotide-binding domain | scaffold_0043 | 34347 | 36920 | -7.0 |
| PCAU.43c3d.1.P00430032 | Aminotransferase, class IV | scaffold_0043 | 41084 | 42443 | -7.3 |
| PCAU.43c3d.1.P00430044 | P-loop containing nucleoside triphosphate hydrolase | scaffold_0043 | 57975 | 58689 | -6.6 |
| PCAU.43c3d.1.P00430061 | Uracil-DNA glycosylase | scaffold_0043 | 77336 | 78201 | -7.1 |
| PCAU.43c3d.1.P00430066 | Protein kinase-like domain | scaffold_0043 | 83338 | 84487 | -7.3 |
| PCAU.43c3d.1.P00430080 | Ribosomal protein S8 | scaffold_0043 | 106504 | 106983 | -5.7 |
| PCAU.43c3d.1.P00430096 | Metallo-dependent phosphatase-like | scaffold_0043 | 133140 | 134863 | -6.4 |
| PCAU.43c3d.1.P00430117 | tRNA-guanine(15) transglycosylase-like | scaffold_0043 | 177075 | 178318 | -7.8 |
| PCAU.43c3d.1.P00430120 | Ubiquitin/SUMO-activating enzyme E1 | scaffold_0043 | 181359 | 184625 | -7.1 |
| PCAU.43c3d.1.P00430133 | Aspartic peptidase | scaffold_0043 | 197365 | 198757 | -8.1 |
| PCAU.43c3d.1.P00430137 | HIT-like domain | scaffold_0043 | 201317 | 201690 | -6.7 |
| PCAU.43c3d.1.P00430139 | HIT-like domain | scaffold_0043 | 202947 | 203425 | -7.3 |
| PCAU.43c3d.1.P00430152 | Ribosomal protein L13e | scaffold_0043 | 238711 | 239447 | -6.3 |
| PCAU.43c3d.1.P00430157 | small GTPase Rab1 family profile. | scaffold_0043 | 245735 | 246446 | -5.6 |
| PCAU.43c3d.1.P00440005 | Peptidase family M49 | scaffold_0044 | 8551 | 10768 | -7.4 |
| PCAU.43c3d.1.P00440011 | Ferrochelatase | scaffold_0044 | 25142 | 26308 | -6.9 |
| PCAU.43c3d.1.P00440034 | Guanylate kinase/L-type calcium channel beta subunit | scaffold_0044 | 58632 | 59247 | -7.8 |
| PCAU.43c3d.1.P00440054 | Elongator protein 3/MiaB/NifB | scaffold_0044 | 90684 | 92437 | -6.4 |
| PCAU.43c3d.1.P00440081 | Protein kinase-like domain | scaffold_0044 | 136259 | 137504 | -6.9 |
| PCAU.43c3d.1.P00450007 | Ubiquitin-conjugating enzyme/RWD-like | scaffold_0045 | 9099 | 9599 | -6.4 |
| PCAU.43c3d.1.P00450073 | Metallo-dependent phosphatase-like | scaffold_0045 | 118684 | 119733 | -6.6 |
| PCAU.43c3d.1.P00450077 | Actin-related protein 4 (Arp4) | scaffold_0045 | 126300 | 127391 | -7.3 |
| PCAU.43c3d.1.P00450078 | Inositol monophosphatase | scaffold_0045 | 127397 | 128419 | -7.5 |
| PCAU.43c3d.1.P00450089 | Aconitase/2-methylisocitrate dehydratase | scaffold_0045 | 148135 | 151010 | -6.9 |
| PCAU.43c3d.1.P00450096 | JAB1/MPN/MOV34 metalloenzyme domain | scaffold_0045 | 159140 | 160123 | -6.9 |
| PCAU.43c3d.1.P00450098 | WD40-repeat-containing domain | scaffold_0045 | 160654 | 162137 | -7.0 |
| PCAU.43c3d.1.P00450136 | EF-hand domain pair | scaffold_0045 | 228317 | 228865 | -7.0 |
| PCAU.43c3d.1.P00450140 | Ribosomal protein L34Ae | scaffold_0045 | 232954 | 233360 | -6.0 |
| PCAU.43c3d.1.P00450169 | Protein kinase-like domain | scaffold_0045 | 280693 | 281771 | -7.4 |
| PCAU.43c3d.1.P00450170 | P-loop containing nucleoside triphosphate hydrolase | scaffold_0045 | 281754 | 283633 | -8.0 |
| PCAU.43c3d.1.P00460074 | Aldehyde/histidinol dehydrogenase | scaffold_0046 | 109573 | 111256 | -7.0 |
| PCAU.43c3d.1.P00460094 | Glycosyl transferase, family 28, C-terminal | scaffold_0046 | 135150 | 135676 | -6.3 |
| PCAU.43c3d.1.P00460161 | Homogentisate 1,2-dioxygenase | scaffold_0046 | 229704 | 231193 | -7.3 |
| PCAU.43c3d.1.P00460166 | Ribosomal protein S4/S9 | scaffold_0046 | 236076 | 236751 | -5.3 |
| PCAU.43c3d.1.P00470024 | DNA replication licensing factor Mcm | scaffold_0047 | 30812 | 33077 | -6.5 |
| PCAU.43c3d.1.P00470036 | NADH:cytochrome b5 reductase (CBR) | scaffold_0047 | 53185 | 54106 | -6.5 |
| PCAU.43c3d.1.P00470047 | Protein kinase domain | scaffold_0047 | 64556 | 65577 | -6.8 |
| PCAU.43c3d.1.P00470067 | Eukaryotic initiation factor 3, gamma subunit | scaffold_0047 | 87315 | 88479 | -6.7 |
| PCAU.43c3d.1.P00470113 | Protein kinase-like domain | scaffold_0047 | 149094 | 150461 | -8.2 |
| PCAU.43c3d.1.P00470150 | Acyl-CoA dehydrogenase/oxidase, N-terminal and middle domain | scaffold_0047 | 201939 | 203213 | -6.5 |
| PCAU.43c3d.1.P00470184 | Actin-related protein | scaffold_0047 | 249713 | 250983 | -6.3 |
| PCAU.43c3d.1.P00470193 | Histone deacetylase superfamily | scaffold_0047 | 260602 | 262094 | -7.3 |
| PCAU.43c3d.1.P00470198 | NAD(P)-binding domain | scaffold_0047 | 268507 | 269435 | -6.7 |
| PCAU.43c3d.1.P00480051 | ATP:guanido phosphotransferase, catalytic domain | scaffold_0048 | 84501 | 85735 | -6.5 |
| PCAU.43c3d.1.P00480071 | ATPase, V1 complex, subunit D | scaffold_0048 | 116053 | 116927 | -6.4 |
| PCAU.43c3d.1.P00480078 | Actin-depolymerising factor homology domain | scaffold_0048 | 129412 | 129939 | -5.4 |
| PCAU.43c3d.1.P00480080 | Nucleotide-binding, alpha-beta plait | scaffold_0048 | 131632 | 132199 | -6.1 |
| PCAU.43c3d.1.P00480153 | Terpenoid cyclases/protein prenyltransferase alpha-alpha toroid | scaffold_0048 | 235471 | 236543 | -7.6 |
| PCAU.43c3d.1.P00480169 | Peptide methionine sulfoxide reductase | scaffold_0048 | 259266 | 259768 | -5.7 |
| PCAU.43c3d.1.P00480170 | Ribosomal protein L21e | scaffold_0048 | 259773 | 260349 | -5.8 |
| PCAU.43c3d.1.P00490021 | WD40-repeat-containing domain | scaffold_0049 | 31356 | 32607 | -6.1 |
| PCAU.43c3d.1.P00500027 | Arsenical pump ATPase, ArsA/GET3 | scaffold_0050 | 54737 | 55828 | -6.7 |
| PCAU.43c3d.1.P00500047 | JmjC domain | scaffold_0050 | 75477 | 76767 | -5.8 |
| PCAU.43c3d.1.P00500060 | Glutathione S-transferase, C-terminal-like | scaffold_0050 | 87982 | 88669 | -6.8 |
| PCAU.43c3d.1.P00500061 | Glutathione S-transferase, C-terminal-like | scaffold_0050 | 89006 | 89780 | -6.1 |
| PCAU.43c3d.1.P00500062 | Glutathione S-transferase, C-terminal-like | scaffold_0050 | 89782 | 90564 | -6.6 |
| PCAU.43c3d.1.P00500063 | Glutathione S-transferase, C-terminal-like | scaffold_0050 | 90634 | 91280 | -6.4 |
| PCAU.43c3d.1.P00500080 | small GTPase Rab1 family profile. | scaffold_0050 | 120769 | 121580 | -7.7 |
| PCAU.43c3d.1.P00500123 | Glutathione S-transferase, C-terminal-like | scaffold_0050 | 218773 | 219460 | -6.9 |
| PCAU.43c3d.1.P00500128 | Glutamine synthetase, catalytic domain | scaffold_0050 | 221985 | 223189 | -6.3 |
| PCAU.43c3d.1.P00500160 | Ribosomal protein L4 domain | scaffold_0050 | 257551 | 258946 | -7.0 |
| PCAU.43c3d.1.P00510005 | P-loop containing nucleoside triphosphate hydrolase | scaffold_0051 | 8133 | 9737 | -7.8 |
| PCAU.43c3d.1.P00510066 | Proteasome component (PCI) domain | scaffold_0051 | 105051 | 106446 | -7.7 |
| PCAU.43c3d.1.P00510079 | Translation initiation factor SUI1 | scaffold_0051 | 124702 | 125112 | -6.3 |
| PCAU.43c3d.1.P00510084 | DNA-directed DNA polymerase, family B | scaffold_0051 | 130594 | 133871 | -6.9 |
| PCAU.43c3d.1.P00510127 | Phosphoglucose isomerase (PGI) | scaffold_0051 | 219198 | 221036 | -6.7 |
| PCAU.43c3d.1.P00510131 | Phosphoribosyltransferase-like | scaffold_0051 | 225149 | 226473 | -6.0 |
| PCAU.43c3d.1.P00510148 | Fructose-bisphosphate aldolase class-I, eukaryotic-type | scaffold_0051 | 243736 | 245013 | -7.7 |
| PCAU.43c3d.1.P00510154 | Ribosomal protein L22/L17, eukaryotic/archaeal | scaffold_0051 | 254964 | 255635 | -5.9 |
| PCAU.43c3d.1.P00510161 | Nucleophile aminohydrolases, N-terminal | scaffold_0051 | 263938 | 264691 | -6.8 |
| PCAU.43c3d.1.P00520010 | P-type ATPase, transmembrane domain | scaffold_0052 | 16841 | 20124 | -7.7 |
| PCAU.43c3d.1.P00520045 | Aminoacyl-tRNA synthetase, class II (D/K/N)-like | scaffold_0052 | 80254 | 82115 | -6.9 |
| PCAU.43c3d.1.P00520088 | Ribosomal protein S21e | scaffold_0052 | 162231 | 162584 | -5.7 |
| PCAU.43c3d.1.P00520106 | Metallo-dependent phosphatase-like | scaffold_0052 | 184993 | 186117 | -6.0 |
| PCAU.43c3d.1.P00520116 | Ribonuclease T2-like | scaffold_0052 | 201431 | 202206 | -5.8 |
| PCAU.43c3d.1.P00520121 | Serine/threonine protein phosphatase 5 | scaffold_0052 | 207202 | 208839 | -6.8 |
| PCAU.43c3d.1.P00520128 | Alanyl-tRNA synthetase, class IIc, core domain | scaffold_0052 | 219366 | 222357 | -7.4 |
| PCAU.43c3d.1.P00520149 | Cyclic nucleotide-binding-like | scaffold_0052 | 256084 | 257234 | -5.8 |
| PCAU.43c3d.1.P00530015 | Calcium/calmodulin-dependent/calcium-dependent protein kinase | scaffold_0053 | 22419 | 23570 | -7.1 |
| PCAU.43c3d.1.P00530034 | Peptidase M1, alanine aminopeptidase/leukotriene A4 hydrolase | scaffold_0053 | 49463 | 51536 | -7.6 |
| PCAU.43c3d.1.P00530057 | Ribosomal protein L24e-related | scaffold_0053 | 78313 | 78899 | -6.9 |
| PCAU.43c3d.1.P00530071 | Small GTPase superfamily, ARF type | scaffold_0053 | 108123 | 108765 | -5.9 |
| PCAU.43c3d.1.P00530108 | Thiolase-like, subgroup | scaffold_0053 | 158305 | 159573 | -7.8 |
| PCAU.43c3d.1.P00530125 | Cyclophilin-like domain | scaffold_0053 | 180438 | 181077 | -8.7 |
| PCAU.43c3d.1.P00530165 | P-type ATPase, transmembrane domain | scaffold_0053 | 239698 | 242958 | -8.0 |
| PCAU.43c3d.1.P00540009 | Cytidine deaminase-like | scaffold_0054 | 10205 | 10749 | -6.1 |
| PCAU.43c3d.1.P00540013 | Pyridoxal phosphate-dependent transferase | scaffold_0054 | 13130 | 14738 | -6.9 |
| PCAU.43c3d.1.P00540021 | DNA-directed RNA polymerase, subunit 2 | scaffold_0054 | 24504 | 27933 | -6.9 |

| | | | | | |
|---|---|---|---|---|---|
| PCAU.43c3d.1.P00540087 | Tubulin | scaffold_0054 | 149752 | 151209 | -6.3 |
| PCAU.43c3d.1.P00540094 | Ribosomal protein L28e | scaffold_0054 | 159502 | 160076 | -6.3 |
| PCAU.43c3d.1.P00540128 | Peptidase M24A, methionine aminopeptidase, subfamily 2 | scaffold_0054 | 209484 | 210968 | -6.0 |
| PCAU.43c3d.1.P00550036 | Autophagy-related protein 3, N-terminal | scaffold_0055 | 48614 | 49428 | -6.5 |
| PCAU.43c3d.1.P00550107 | Ribosomal protein S3, C-terminal | scaffold_0055 | 157446 | 158264 | -6.3 |
| PCAU.43c3d.1.P00550116 | Glutaredoxin | scaffold_0055 | 170881 | 171211 | -5.3 |
| PCAU.43c3d.1.P00560001 | CoA-transferase family III | scaffold_0056 | 360 | 1480 | -7.2 |
| PCAU.43c3d.1.P00560011 | Ribosomal protein L14b/L23e | scaffold_0056 | 14885 | 15404 | -6.5 |
| PCAU.43c3d.1.P00560103 | EF-hand domain | scaffold_0056 | 131879 | 132430 | -6.1 |
| PCAU.43c3d.1.P00560143 | Plectin/S10, N-terminal | scaffold_0056 | 219194 | 219719 | -7.0 |
| PCAU.43c3d.1.P00560147 | Elongation factor G, III-V domain | scaffold_0056 | 220999 | 223187 | -6.7 |
| PCAU.43c3d.1.P00560148 | P-loop containing nucleoside triphosphate hydrolase | scaffold_0056 | 223221 | 223952 | -6.1 |
| PCAU.43c3d.1.P00560157 | Isocitrate and isopropylmalate dehydrogenases family | scaffold_0056 | 233027 | 234200 | -6.8 |
| PCAU.43c3d.1.P00570027 | small GTPase Rab1 family profile. | scaffold_0057 | 38211 | 38861 | -6.6 |
| PCAU.43c3d.1.P00570032 | Protein kinase-like domain | scaffold_0057 | 53092 | 54172 | -7.3 |
| PCAU.43c3d.1.P00570036 | Metallo-dependent phosphatase-like | scaffold_0057 | 58204 | 59324 | -6.0 |
| PCAU.43c3d.1.P00570102 | ATP:guanido phosphotransferase, catalytic domain | scaffold_0057 | 161302 | 162507 | -7.0 |
| PCAU.43c3d.1.P00570145 | Ribosomal protein S24e | scaffold_0057 | 248342 | 248934 | -6.8 |
| PCAU.43c3d.1.P00580001 | small GTPase Rab1 family profile. | scaffold_0058 | 1051 | 1805 | -6.8 |
| PCAU.43c3d.1.P00580006 | Uroporphyrinogen decarboxylase (URO-D) | scaffold_0058 | 6046 | 7350 | -7.1 |
| PCAU.43c3d.1.P00580035 | V-ATPase proteolipid subunit C-like domain | scaffold_0058 | 52462 | 53063 | -5.7 |
| PCAU.43c3d.1.P00580040 | Histone deacetylase superfamily | scaffold_0058 | 56617 | 58150 | -6.5 |
| PCAU.43c3d.1.P00580063 | Serine-tRNA ligase, type1 | scaffold_0058 | 85794 | 87311 | -6.9 |
| PCAU.43c3d.1.P00580086 | Zinc finger, ZPR1-type | scaffold_0058 | 114363 | 115870 | -6.4 |
| PCAU.43c3d.1.P00580087 | Zinc finger, ZPR1-type | scaffold_0058 | 115974 | 116789 | -6.7 |
| PCAU.43c3d.1.P00580090 | Longin-like domain | scaffold_0058 | 118465 | 119170 | -6.4 |
| PCAU.43c3d.1.P00580103 | Ribosomal protein L7A/L8 | scaffold_0058 | 142530 | 143446 | -6.4 |
| PCAU.43c3d.1.P00580163 | Aldolase-type TIM barrel | scaffold_0058 | 235957 | 237490 | -6.9 |
| PCAU.43c3d.1.P00580166 | Protein kinase-like domain | scaffold_0058 | 241866 | 243151 | -7.7 |
| PCAU.43c3d.1.P00590040 | Sugar-phosphate isomerase, RpiB/LacA/LacB family | scaffold_0059 | 49759 | 50178 | -6.7 |
| PCAU.43c3d.1.P00590046 | Peptidase C26, gamma-glutamyl hydrolase | scaffold_0059 | 58502 | 59527 | -7.8 |
| PCAU.43c3d.1.P00590087 | Thioredoxin | scaffold_0059 | 101535 | 101977 | -5.7 |
| PCAU.43c3d.1.P00590108 | RmlC-like jelly roll fold | scaffold_0059 | 140759 | 141921 | -6.3 |
| PCAU.43c3d.1.P00590128 | P-loop containing nucleoside triphosphate hydrolase | scaffold_0059 | 174560 | 175288 | -7.2 |
| PCAU.43c3d.1.P00590136 | Pyruvate dehydrogenase E1 component subunit beta | scaffold_0059 | 185876 | 187009 | -6.3 |
| PCAU.43c3d.1.P00590144 | mRNA (2'-O-methyladenosine-N(6)-)-methyltransferase-like (MT-A70-like) family profile. | scaffold_0059 | 193647 | 194698 | -7.2 |
| PCAU.43c3d.1.P00590172 | Protein kinase-like domain | scaffold_0059 | 241112 | 242137 | -6.4 |
| PCAU.43c3d.1.P00600061 | Protein kinase-like domain | scaffold_0060 | 100958 | 102095 | -6.5 |
| PCAU.43c3d.1.P00600082 | Glutathione S-transferase, C-terminal-like | scaffold_0060 | 142366 | 143111 | -6.9 |
| PCAU.43c3d.1.P00600110 | Pyruvate/Phosphoenolpyruvate kinase-like domain | scaffold_0060 | 193098 | 194069 | -6.4 |
| PCAU.43c3d.1.P00600133 | Ribosomal protein S2, eukaryotic/archaeal | scaffold_0060 | 227184 | 228074 | -6.0 |
| PCAU.43c3d.1.P00610106 | Fumarylacetoacetase | scaffold_0061 | 184574 | 186007 | -6.8 |
| PCAU.43c3d.1.P00610113 | Peptidase family M49 | scaffold_0061 | 196069 | 198193 | -7.7 |
| PCAU.43c3d.1.P00610130 | Janus | scaffold_0061 | 222770 | 223250 | -5.5 |
| PCAU.43c3d.1.P00620009 | Protein kinase-like domain | scaffold_0062 | 18556 | 19533 | -6.6 |
| PCAU.43c3d.1.P00620025 | ABC transporter A, ABCA | scaffold_0062 | 34828 | 38909 | -7.6 |
| PCAU.43c3d.1.P00620033 | Tetratricopeptide repeat | scaffold_0062 | 51106 | 52799 | -6.7 |
| PCAU.43c3d.1.P00620039 | Cyclin-dependent kinase, regulatory subunit | scaffold_0062 | 58174 | 58439 | -6.1 |
| PCAU.43c3d.1.P00620058 | Peptide chain release factor eRF1/aRF1 | scaffold_0062 | 83692 | 84227 | -6.9 |
| PCAU.43c3d.1.P00620128 | Phenylalanyl-tRNA synthetase | scaffold_0062 | 173017 | 174228 | -8.2 |
| PCAU.43c3d.1.P00630029 | Protein translocase SecE domain | scaffold_0063 | 43111 | 43392 | -5.2 |
| PCAU.43c3d.1.P00630054 | Alpha-helical ferredoxin | scaffold_0063 | 94116 | 95131 | -6.6 |
| PCAU.43c3d.1.P00630067 | Nucleophile aminohydrolases, N-terminal | scaffold_0063 | 115474 | 116269 | -6.3 |
| PCAU.43c3d.1.P00630088 | Mannose-6-phosphate isomerase | scaffold_0063 | 143988 | 144832 | -6.7 |
| PCAU.43c3d.1.P00630089 | Mannose-6-phosphate isomerase | scaffold_0063 | 144641 | 145185 | -6.3 |
| PCAU.43c3d.1.P00630130 | P-loop containing nucleoside triphosphate hydrolase | scaffold_0063 | 215814 | 216488 | -6.1 |
| PCAU.43c3d.1.P00640007 | Peptidyl-prolyl cis-trans isomerase, FKBP-type | scaffold_0064 | 9175 | 9535 | -5.9 |
| PCAU.43c3d.1.P00640017 | Myo-inositol-1-phosphate synthase | scaffold_0064 | 29381 | 30961 | -7.4 |
| PCAU.43c3d.1.P00640028 | Glycoside hydrolase, family 37 | scaffold_0064 | 40787 | 42433 | -6.4 |
| PCAU.43c3d.1.P00640054 | Glutamine synthetase/guanido kinase, catalytic domain | scaffold_0064 | 82832 | 83989 | -7.5 |
| PCAU.43c3d.1.P00640063 | DNA recombination and repair protein Rad51, C-terminal | scaffold_0064 | 94051 | 95089 | -6.4 |
| PCAU.43c3d.1.P00640066 | P-loop containing nucleoside triphosphate hydrolase | scaffold_0064 | 97706 | 98983 | -6.8 |
| PCAU.43c3d.1.P00640148 | Thiamin diphosphate-binding fold | scaffold_0064 | 219425 | 221564 | -6.7 |
| PCAU.43c3d.1.P00650015 | P-loop containing nucleoside triphosphate hydrolase | scaffold_0065 | 23691 | 24475 | -6.8 |
| PCAU.43c3d.1.P00650064 | Zinc finger, RING/FYVE/PHD-type | scaffold_0065 | 110903 | 111276 | -5.5 |
| PCAU.43c3d.1.P00650080 | Thioredoxin-like fold | scaffold_0065 | 132995 | 133856 | -5.8 |
| PCAU.43c3d.1.P00650111 | Metallo-dependent phosphatase-like | scaffold_0065 | 175673 | 177172 | -6.5 |
| PCAU.43c3d.1.P00650113 | Acyl-CoA dehydrogenase/oxidase C-terminal | scaffold_0065 | 178711 | 180766 | -7.4 |
| PCAU.43c3d.1.P00650137 | Cyclic nucleotide-binding domain | scaffold_0065 | 210132 | 210971 | -7.9 |
| PCAU.43c3d.1.P00660090 | P-loop containing nucleoside triphosphate hydrolase | scaffold_0066 | 137238 | 138610 | -7.5 |
| PCAU.43c3d.1.P00660099 | Arginine-tRNA ligase | scaffold_0066 | 149084 | 150948 | -6.7 |
| PCAU.43c3d.1.P00660120 | NMD3 | scaffold_0066 | 179751 | 181150 | -6.7 |
| PCAU.43c3d.1.P00660122 | Protein kinase-like domain | scaffold_0066 | 182706 | 184107 | -7.0 |
| PCAU.43c3d.1.P00670016 | Peptidase M16, C-terminal domain | scaffold_0067 | 20945 | 23932 | -7.5 |
| PCAU.43c3d.1.P00670025 | Saccharopine dehydrogenase / Homospermidine synthase | scaffold_0067 | 33332 | 34891 | -7.1 |
| PCAU.43c3d.1.P00670029 | Small GTPase superfamily, ARF type | scaffold_0067 | 39201 | 39872 | -5.9 |
| PCAU.43c3d.1.P00670048 | Zinc finger, RING/FYVE/PHD-type | scaffold_0067 | 65456 | 65831 | -5.3 |
| PCAU.43c3d.1.P00670050 | Ornithine aminotransferase | scaffold_0067 | 68031 | 69426 | -6.8 |
| PCAU.43c3d.1.P00670055 | Protein kinase-like domain | scaffold_0067 | 76741 | 79367 | -7.7 |
| PCAU.43c3d.1.P00670081 | Pyridoxal phosphate-dependent transferase | scaffold_0067 | 113530 | 114897 | -7.6 |
| PCAU.43c3d.1.P00670091 | Ubiquitin-conjugating enzyme/RWD-like | scaffold_0067 | 145761 | 146463 | -6.7 |
| PCAU.43c3d.1.P00670092 | P-loop containing nucleoside triphosphate hydrolase | scaffold_0067 | 146457 | 147879 | -6.0 |
| PCAU.43c3d.1.P00670098 | Ribosomal RNA assembly KRR1 | scaffold_0067 | 155795 | 156833 | -6.2 |
| PCAU.43c3d.1.P00670106 | Gcp-like domain | scaffold_0067 | 167935 | 169217 | -7.3 |
| PCAU.43c3d.1.P00670122 | small GTPase Rab1 family profile. | scaffold_0067 | 193272 | 194011 | -5.9 |
| PCAU.43c3d.1.P00680002 | Protein kinase domain | scaffold_0068 | 2428 | 3563 | -7.3 |
| PCAU.43c3d.1.P00680028 | Aromatic amino acid hydroxylase | scaffold_0068 | 48615 | 49996 | -8.9 |
| PCAU.43c3d.1.P00680082 | Ribosomal protein L30, ferredoxin-like fold domain | scaffold_0068 | 139867 | 140732 | -6.6 |

| | | | | | |
|---|---|---|---|---|---|
| PCAU.43c3d.1.P00680119 | P-loop containing nucleoside triphosphate hydrolase | scaffold_0068 | 218511 | 219217 | -8.5 |
| PCAU.43c3d.1.P00690021 | ABC transporter, transmembrane domain | scaffold_0069 | 29407 | 33445 | -7.3 |
| PCAU.43c3d.1.P00690108 | Alpha-D-phosphohexomutase, alpha/beta/alpha I/II/III | scaffold_0069 | 145144 | 146706 | -6.8 |
| PCAU.43c3d.1.P00700035 | P-loop containing nucleoside triphosphate hydrolase | scaffold_0070 | 49552 | 50241 | -8.0 |
| PCAU.43c3d.1.P00700137 | Protein farnesyltransferase subunit beta | scaffold_0070 | 185805 | 186982 | -6.1 |
| PCAU.43c3d.1.P00710010 | Phytanoyl-CoA dioxygenase | scaffold_0071 | 20195 | 21249 | -7.3 |
| PCAU.43c3d.1.P00710027 | Ribosomal protein S8e | scaffold_0071 | 44091 | 44795 | -6.0 |
| PCAU.43c3d.1.P00710065 | 2-oxoglutarate dehydrogenase E1 component | scaffold_0071 | 92751 | 95793 | -6.9 |
| PCAU.43c3d.1.P00710068 | EF-hand domain pair | scaffold_0071 | 98601 | 99101 | -6.0 |
| PCAU.43c3d.1.P00710074 | Protein kinase-like domain | scaffold_0071 | 105484 | 106374 | -6.2 |
| PCAU.43c3d.1.P00710081 | UDP-glucose 4-epimerase GalE | scaffold_0071 | 114003 | 115043 | -8.3 |
| PCAU.43c3d.1.P00720003 | Acyl-CoA dehydrogenase/oxidase C-terminal | scaffold_0072 | 1356 | 3464 | -7.2 |
| PCAU.43c3d.1.P00720055 | Adenosine kinase | scaffold_0072 | 85811 | 86939 | -7.1 |
| PCAU.43c3d.1.P00720074 | Nucleophile aminohydrolases, N-terminal | scaffold_0072 | 117810 | 118648 | -6.5 |
| PCAU.43c3d.1.P00720091 | Translation initiation factor IF6 | scaffold_0072 | 140250 | 141121 | -5.7 |
| PCAU.43c3d.1.P00720103 | NAD(P)-binding domain | scaffold_0072 | 163462 | 164415 | -7.0 |
| PCAU.43c3d.1.P00720104 | small GTPase Rab1 family profile. | scaffold_0072 | 164450 | 165189 | -6.2 |
| PCAU.43c3d.1.P00720129 | Superoxide dismutase, copper/zinc binding domain | scaffold_0072 | 193868 | 194517 | -6.3 |
| PCAU.43c3d.1.P00730032 | Aspartic peptidase | scaffold_0073 | 45256 | 46660 | -6.7 |
| PCAU.43c3d.1.P00730033 | Metallo-dependent phosphatase-like | scaffold_0073 | 46673 | 48214 | -6.2 |
| PCAU.43c3d.1.P00730056 | ClpP/crotonase-like domain | scaffold_0073 | 70845 | 71743 | -6.6 |
| PCAU.43c3d.1.P00730062 | Cyclin-dependent kinase, regulatory subunit | scaffold_0073 | 81552 | 81916 | -7.1 |
| PCAU.43c3d.1.P00730125 | Phosphofructokinase domain | scaffold_0073 | 185701 | 187445 | -6.8 |
| PCAU.43c3d.1.P00730136 | Tubulin | scaffold_0073 | 200062 | 201452 | -6.8 |
| PCAU.43c3d.1.P00740012 | V-ATPase proteolipid subunit | scaffold_0074 | 22321 | 23011 | -6.0 |
| PCAU.43c3d.1.P00740034 | Cyclophilin-like domain | scaffold_0074 | 47947 | 48567 | -6.7 |
| PCAU.43c3d.1.P00740050 | Glutathione S-transferase, C-terminal-like | scaffold_0074 | 79841 | 80541 | -6.5 |
| PCAU.43c3d.1.P00740051 | Coiled coil domain | scaffold_0074 | 80575 | 82003 | -6.5 |
| PCAU.43c3d.1.P00740055 | Glutathione S-transferase, C-terminal-like | scaffold_0074 | 84745 | 85489 | -6.8 |
| PCAU.43c3d.1.P00740067 | Malate dehydrogenase, type 2 | scaffold_0074 | 110067 | 111252 | -6.3 |
| PCAU.43c3d.1.P00740068 | NUDIX hydrolase domain-like | scaffold_0074 | 111275 | 111868 | -6.1 |
| PCAU.43c3d.1.P00740111 | Ureohydrolase | scaffold_0074 | 169592 | 170584 | -6.8 |
| PCAU.43c3d.1.P00740127 | Glycosyl transferase, family 4 | scaffold_0074 | 188903 | 190078 | -7.6 |
| PCAU.43c3d.1.P00750013 | Aldehyde/histidinol dehydrogenase | scaffold_0075 | 14123 | 15813 | -7.2 |
| PCAU.43c3d.1.P00750047 | Glycoside hydrolase, family 13 | scaffold_0075 | 69764 | 71286 | -6.5 |
| PCAU.43c3d.1.P00750056 | Phosphoribosyltransferase-like | scaffold_0075 | 85029 | 85906 | -6.4 |
| PCAU.43c3d.1.P00750060 | Ubiquitin-like modifier-activating enzyme Atg7 | scaffold_0075 | 89813 | 91873 | -7.1 |
| PCAU.43c3d.1.P00750068 | small GTPase Rab1 family profile. | scaffold_0075 | 100540 | 101277 | -6.4 |
| PCAU.43c3d.1.P00750076 | Actin-related protein | scaffold_0075 | 114945 | 116249 | -6.7 |
| PCAU.43c3d.1.P00750104 | Calcium/calmodulin-dependent/calcium-dependent protein kinase | scaffold_0075 | 149912 | 150893 | -6.6 |
| PCAU.43c3d.1.P00760002 | Acyl-CoA dehydrogenase/oxidase, N-terminal and middle domain | scaffold_0076 | 3850 | 5929 | -7.6 |
| PCAU.43c3d.1.P00760032 | Small GTPase superfamily, ARF type | scaffold_0076 | 44887 | 45517 | -5.6 |
| PCAU.43c3d.1.P00760095 | Tetratricopeptide repeat | scaffold_0076 | 164864 | 166949 | -6.3 |
| PCAU.43c3d.1.P00760107 | Glutaredoxin | scaffold_0076 | 178425 | 178764 | -7.0 |
| PCAU.43c3d.1.P00760109 | RNA 3'-terminal phosphate cyclase | scaffold_0076 | 179595 | 180700 | -6.1 |
| PCAU.43c3d.1.P00770009 | Citrate synthase-like | scaffold_0077 | 10821 | 12434 | -7.3 |
| PCAU.43c3d.1.P00770133 | Small GTPase superfamily, ARF type | scaffold_0077 | 179481 | 180027 | -6.3 |
| PCAU.43c3d.1.P00780040 | Protein kinase-like domain | scaffold_0078 | 59449 | 60680 | -6.9 |
| PCAU.43c3d.1.P00780090 | small GTPase Rab1 family profile. | scaffold_0078 | 131988 | 132736 | -6.3 |
| PCAU.43c3d.1.P00780095 | Metallo-dependent phosphatase-like | scaffold_0078 | 142010 | 143109 | -6.8 |
| PCAU.43c3d.1.P00790004 | GTP-binding protein Obg/CgtA | scaffold_0079 | 12061 | 13203 | -6.1 |
| PCAU.43c3d.1.P00790021 | Small GTPase superfamily, SAR1-type | scaffold_0079 | 32054 | 32733 | -6.7 |
| PCAU.43c3d.1.P00790057 | Ribosomal protein S17 | scaffold_0079 | 87909 | 88494 | -6.1 |
| PCAU.43c3d.1.P00790080 | Ribosomal protein L15e | scaffold_0079 | 112674 | 113410 | -5.9 |
| PCAU.43c3d.1.P00790085 | Ribosomal protein L39e | scaffold_0079 | 119670 | 119885 | -5.4 |
| PCAU.43c3d.1.P00790097 | Ribosomal protein L2 | scaffold_0079 | 133314 | 134237 | -7.2 |
| PCAU.43c3d.1.P00800040 | Translation Initiation factor eIF- 4e | scaffold_0080 | 65008 | 65703 | -7.1 |
| PCAU.43c3d.1.P00800061 | Coenzyme A transferase family I | scaffold_0080 | 95760 | 97341 | -8.2 |
| PCAU.43c3d.1.P00800078 | Phosphoglycerate kinase | scaffold_0080 | 121783 | 123143 | -6.8 |
| PCAU.43c3d.1.P00800089 | ATPase, F1 complex, gamma subunit | scaffold_0080 | 136015 | 137003 | -6.6 |
| PCAU.43c3d.1.P00800090 | P-loop containing nucleoside triphosphate hydrolase | scaffold_0080 | 137031 | 138571 | -6.7 |
| PCAU.43c3d.1.P00800105 | Adenylate kinase/UMP-CMP kinase | scaffold_0080 | 166334 | 167786 | -6.4 |
| PCAU.43c3d.1.P00800109 | Chaperonin Cpn10 | scaffold_0080 | 171336 | 171719 | -5.8 |
| PCAU.43c3d.1.P00800110 | T-complex protein 1, gamma subunit | scaffold_0080 | 171771 | 173572 | -6.4 |
| PCAU.43c3d.1.P00810010 | Hydroxymethylglutaryl-CoA lyase | scaffold_0081 | 15444 | 16497 | -7.1 |
| PCAU.43c3d.1.P00810024 | Ribosomal protein S5 | scaffold_0081 | 37627 | 38547 | -6.5 |
| PCAU.43c3d.1.P00810065 | Armadillo-type fold | scaffold_0081 | 86789 | 88728 | -6.0 |
| PCAU.43c3d.1.P00810070 | Translation-associated RNA-binding, predicted | scaffold_0081 | 92990 | 93567 | -6.9 |
| PCAU.43c3d.1.P00810092 | Ribosomal protein L13 | scaffold_0081 | 126148 | 126850 | -6.8 |
| PCAU.43c3d.1.P00810101 | Ribosomal protein L44e | scaffold_0081 | 136900 | 137300 | -5.9 |
| PCAU.43c3d.1.P00820032 | Fumarylacetoacetase, C-terminal | scaffold_0082 | 42184 | 42909 | -5.7 |
| PCAU.43c3d.1.P00820035 | Major facilitator superfamily domain, general substrate transporter | scaffold_0082 | 44996 | 46573 | -6.1 |
| PCAU.43c3d.1.P00820046 | CAAX prenyl protease 1 | scaffold_0082 | 63243 | 64727 | -6.7 |
| PCAU.43c3d.1.P00820048 | NAD(P)-binding domain | scaffold_0082 | 65757 | 66668 | -7.6 |
| PCAU.43c3d.1.P00820054 | Citrate synthase-like | scaffold_0082 | 76416 | 77978 | -7.5 |
| PCAU.43c3d.1.P00820096 | P-loop containing nucleoside triphosphate hydrolase | scaffold_0082 | 131025 | 131745 | -7.2 |
| PCAU.43c3d.1.P00820108 | Protein phosphatase 2C (PP2C)-like domain | scaffold_0082 | 142755 | 143806 | -6.7 |
| PCAU.43c3d.1.P00820118 | Mitochondrial substrate/solute carrier | scaffold_0082 | 157529 | 158614 | -7.9 |
| PCAU.43c3d.1.P00820138 | Fructose-1,6-bisphosphatase class 1/Sedoheputulose-1,7-bisphosphatase | scaffold_0082 | 178202 | 179276 | -6.3 |
| PCAU.43c3d.1.P00830071 | Protein kinase-like domain | scaffold_0083 | 93186 | 94279 | -7.0 |
| PCAU.43c3d.1.P00840015 | small GTPase Rab1 family profile. | scaffold_0084 | 16811 | 17493 | -7.5 |
| PCAU.43c3d.1.P00840049 | P-loop containing nucleoside triphosphate hydrolase | scaffold_0084 | 62114 | 63876 | -6.6 |
| PCAU.43c3d.1.P00840056 | Peptide methionine sulphoxide reductase MsrA | scaffold_0084 | 72226 | 72759 | -5.9 |
| PCAU.43c3d.1.P00840097 | Mini-chromosome maintenance, DNA-dependent ATPase | scaffold_0084 | 123033 | 125483 | -7.2 |
| PCAU.43c3d.1.P00850005 | NAD(P)-binding domain | scaffold_0085 | 27092 | 28825 | -7.2 |
| PCAU.43c3d.1.P00850012 | 26S proteasome, regulatory subunit Rpn7 | scaffold_0085 | 41415 | 42468 | -7.7 |
| PCAU.43c3d.1.P00850051 | Pyridoxal phosphate-dependent transferase | scaffold_0085 | 88019 | 89351 | -6.3 |
| PCAU.43c3d.1.P00850096 | Citrate synthase-like | scaffold_0085 | 159015 | 160552 | -6.3 |

| | | | | | |
|---|---|---|---|---|---|
| PCAU.43c3d.1.P00860017 | NAD(P)-binding domain | scaffold_0086 | 20860 | 22496 | -6.7 |
| PCAU.43c3d.1.P00860061 | Thioredoxin-like fold | scaffold_0086 | 86007 | 86441 | -6.2 |
| PCAU.43c3d.1.P00860062 | tRNA (guanine-N-7) methyltransferase catalytic subunit Trm8, eukaryote | scaffold_0086 | 86508 | 87354 | -7.9 |
| PCAU.43c3d.1.P00860070 | Glyceraldehyde/Erythrose phosphate dehydrogenase family | scaffold_0086 | 93980 | 95186 | -6.3 |
| PCAU.43c3d.1.P00860072 | Ribosomal protein S28e | scaffold_0086 | 96884 | 97112 | -6.1 |
| PCAU.43c3d.1.P00860081 | Heat shock protein 70 family | scaffold_0086 | 114623 | 116649 | -7.3 |
| PCAU.43c3d.1.P00860083 | Cyclic nucleotide-binding-like | scaffold_0086 | 117521 | 119218 | -6.6 |
| PCAU.43c3d.1.P00860114 | Thiamin diphosphate-binding fold | scaffold_0086 | 159306 | 160496 | -7.4 |
| PCAU.43c3d.1.P00870008 | Glycerol-3-phosphate dehydrogenase, NAD-dependent | scaffold_0087 | 11389 | 12492 | -7.9 |
| PCAU.43c3d.1.P00870013 | Polyprenyl synthetase-related | scaffold_0087 | 17582 | 18723 | -8.1 |
| PCAU.43c3d.1.P00870017 | Polyprenyl synthetase-related | scaffold_0087 | 19850 | 21061 | -6.9 |
| PCAU.43c3d.1.P00870034 | Porphobilinogen synthase | scaffold_0087 | 43413 | 44429 | -6.1 |
| PCAU.43c3d.1.P00880001 | Isopentenyl-diphosphate delta-isomerase, type 1 | scaffold_0088 | 238 | 1030 | -5.5 |
| PCAU.43c3d.1.P00880008 | small GTPase Rab1 family profile. | scaffold_0088 | 13294 | 13975 | -6.4 |
| PCAU.43c3d.1.P00880013 | Ribosomal protein S19/S15 | scaffold_0088 | 21054 | 21612 | -5.6 |
| PCAU.43c3d.1.P00880018 | EF-hand domain | scaffold_0088 | 26388 | 26941 | -7.0 |
| PCAU.43c3d.1.P00880024 | Protein kinase domain | scaffold_0088 | 42083 | 43304 | -6.3 |
| PCAU.43c3d.1.P00880027 | Isocitrate dehydrogenase NADP-dependent | scaffold_0088 | 46686 | 48054 | -6.8 |
| PCAU.43c3d.1.P00880059 | P-loop containing nucleoside triphosphate hydrolase | scaffold_0088 | 92112 | 92802 | -6.8 |
| PCAU.43c3d.1.P00880067 | ABC transporter-like | scaffold_0088 | 101750 | 102754 | -6.7 |
| PCAU.43c3d.1.P00880091 | Protein kinase-like domain | scaffold_0088 | 143985 | 145067 | -7.4 |
| PCAU.43c3d.1.P00890077 | NAD(P)-binding domain | scaffold_0089 | 119335 | 120070 | -6.9 |
| PCAU.43c3d.1.P00900005 | Heat shock protein 70 family | scaffold_0090 | 7481 | 9626 | -7.7 |
| PCAU.43c3d.1.P00900006 | Glycoside hydrolase, family 31 | scaffold_0090 | 9594 | 12243 | -6.7 |
| PCAU.43c3d.1.P00900015 | Transaldolase | scaffold_0090 | 19049 | 20310 | -6.2 |
| PCAU.43c3d.1.P00900027 | Mini-chromosome maintenance, DNA-dependent ATPase | scaffold_0090 | 35462 | 38127 | -7.5 |
| PCAU.43c3d.1.P00900029 | Acyl-CoA dehydrogenase/oxidase C-terminal | scaffold_0090 | 38492 | 40543 | -7.1 |
| PCAU.43c3d.1.P00900068 | small GTPase Rab1 family profile. | scaffold_0090 | 100538 | 101331 | -6.8 |
| PCAU.43c3d.1.P00900115 | Oligosaccharyl transferase, STT3 subunit | scaffold_0090 | 164801 | 167104 | -7.2 |
| PCAU.43c3d.1.P00910001 | Translation Initiation factor eIF- 4e-like domain | scaffold_0091 | 483 | 1068 | -6.3 |
| PCAU.43c3d.1.P00910039 | Small GTPase superfamily, ARF type | scaffold_0091 | 56967 | 57574 | -6.0 |
| PCAU.43c3d.1.P00910054 | Alcohol dehydrogenase superfamily, zinc-type | scaffold_0091 | 77890 | 79055 | -7.4 |
| PCAU.43c3d.1.P00910085 | V-type ATPase, V0 complex, 116kDa subunit family | scaffold_0091 | 115439 | 117985 | -7.1 |
| PCAU.43c3d.1.P00910095 | P-loop containing nucleoside triphosphate hydrolase | scaffold_0091 | 128219 | 128827 | -6.9 |
| PCAU.43c3d.1.P00910120 | Protein-tyrosine phosphatase-like | scaffold_0091 | 165901 | 167319 | -6.2 |
| PCAU.43c3d.1.P00920008 | P-loop containing nucleoside triphosphate hydrolase | scaffold_0092 | 12971 | 14471 | -5.9 |
| PCAU.43c3d.1.P00920018 | P-loop containing nucleoside triphosphate hydrolase | scaffold_0092 | 30666 | 31416 | -7.8 |
| PCAU.43c3d.1.P00920037 | Ribosomal protein L6 | scaffold_0092 | 62924 | 63617 | -5.5 |
| PCAU.43c3d.1.P00920038 | Rossmann-like alpha/beta/alpha sandwich fold | scaffold_0092 | 63575 | 64961 | -6.7 |
| PCAU.43c3d.1.P00920045 | Ribosomal protein S24e | scaffold_0092 | 72014 | 72622 | -5.8 |
| PCAU.43c3d.1.P00920074 | P-loop containing nucleoside triphosphate hydrolase | scaffold_0092 | 123919 | 124852 | -6.1 |
| PCAU.43c3d.1.P00920075 | Protein-L-isoaspartate(D-aspartate) O-methyltransferase | scaffold_0092 | 124861 | 125575 | -5.9 |
| PCAU.43c3d.1.P00930020 | Zinc finger, TFIIS-type | scaffold_0093 | 28056 | 28370 | -5.9 |
| PCAU.43c3d.1.P00930049 | P-loop containing nucleoside triphosphate hydrolase | scaffold_0093 | 60224 | 61049 | -8.0 |
| PCAU.43c3d.1.P00930072 | Ribosomal protein L5 | scaffold_0093 | 96203 | 96893 | -6.9 |
| PCAU.43c3d.1.P00930073 | Ribosomal protein S3Ae | scaffold_0093 | 96903 | 97867 | -6.6 |
| PCAU.43c3d.1.P00930075 | Flavoprotein pyridine nucleotide cytochrome reductase | scaffold_0093 | 98720 | 100561 | -7.9 |
| PCAU.43c3d.1.P00930086 | Trimeric LpxA-like | scaffold_0093 | 110914 | 111619 | -7.1 |
| PCAU.43c3d.1.P00930106 | Cyclic nucleotide-binding-like | scaffold_0093 | 147020 | 149658 | -7.9 |
| PCAU.43c3d.1.P00940009 | Pyridoxal phosphate-dependent transferase | scaffold_0094 | 21200 | 22395 | -7.4 |
| PCAU.43c3d.1.P00940016 | Metallo-dependent phosphatase-like | scaffold_0094 | 28284 | 29419 | -6.4 |
| PCAU.43c3d.1.P00940037 | Thioredoxin-like fold | scaffold_0094 | 68159 | 68555 | -6.1 |
| PCAU.43c3d.1.P00940069 | Small GTPase superfamily, ARF type | scaffold_0094 | 121129 | 121672 | -5.7 |
| PCAU.43c3d.1.P00950001 | Pirin | scaffold_0095 | 520 | 1452 | -7.1 |
| PCAU.43c3d.1.P00950041 | Protein kinase-like domain | scaffold_0095 | 55754 | 56954 | -8.6 |
| PCAU.43c3d.1.P00950066 | tRNA methyltransferase, Trm1 | scaffold_0095 | 93666 | 94755 | -6.2 |
| PCAU.43c3d.1.P00950073 | small GTPase Rab1 family profile. | scaffold_0095 | 102811 | 103487 | -6.6 |
| PCAU.43c3d.1.P00950075 | ARP2/3 complex, 20kDa subunit (P20-Arc) | scaffold_0095 | 104992 | 105619 | -6.5 |
| PCAU.43c3d.1.P01260004 | Succinyl-CoA ligase, alpha subunit | scaffold_0126 | 7716 | 8836 | -6.0 |
| PCAU.43c3d.1.P01520001 | Cytochrome c, class IA/ IB | scaffold_0152 | 1160 | 1612 | -6.9 |
| PCAU.43c3d.1.P01630002 | Alpha carbonic anhydrase | scaffold_0163 | 1105 | 1697 | -6.2 |
| PCAU.43c3d.1.P01930003 | Ribosomal protein S5 domain 2-type fold, subgroup | scaffold_0193 | 2706 | 3262 | -5.9 |
| PCAU.43c3d.1.P02300001 | Thiolase-like, subgroup | scaffold_0230 | 279 | 1505 | -6.8 |
| PCAU.43c3d.1.P03640001 | S-adenosyl-L-methionine-dependent methyltransferase-like | scaffold_0364 | 320 | 1168 | -6.7 |

**Table S2.** KEGG pathway categories of the 852 high-quality modelled proteins from *P. caudatum* proteome.

| KEGG pathway | Proteins | KEGG pathway | Proteins |
|---|---|---|---|
| Metabolic pathways | 204 | Biosynthesis of unsaturated fatty acids | 9 |
| Ribosome | 136 | Starch and sucrose metabolism | 9 |
| Biosynthesis of secondary metabolites | 94 | Cysteine and methionine metabolism | 9 |
| Biosynthesis of antibiotics | 80 | Protein processing in endoplasmic reticulum | 18 |
| Carbon metabolism | 58 | Ribosome biogenesis in eukaryotes | 14 |
| Biosynthesis of amino acids | 42 | Ubiquitin mediated proteolysis | 16 |
| Citrate cycle (TCA cycle) | 28 | mRNA surveillance pathway | 12 |
| Purine metabolism | 34 | Nitrogen metabolism | 6 |
| Phagosome | 32 | Terpenoid backbone biosynthesis | 6 |
| Proteasome | 24 | Arachidonic acid metabolism | 6 |
| Glyoxylate and dicarboxylate metabolism | 19 | Butanoate metabolism | 5 |
| Glycolysis / Gluconeogenesis | 19 | beta-Alanine metabolism | 5 |
| Pyrimidine metabolism | 24 | Amino sugar and nucleotide sugar metabolism | 8 |
| Peroxisome | 21 | AGE-RAGE signaling pathway in diabetic complications | 7 |
| Valine, leucine and isoleucine degradation | 16 | Autophagy - other | 11 |
| 2-Oxocarboxylic acid metabolism | 15 | Porphyrin and chlorophyll metabolism | 5 |
| Aminoacyl-tRNA biosynthesis | 16 | Arginine and proline metabolism | 6 |
| Pentose phosphate pathway | 12 | Thiamine metabolism | 6 |
| Oxidative phosphorylation | 22 | Histidine metabolism | 5 |
| Pyruvate metabolism | 13 | Folate biosynthesis | 6 |
| Propanoate metabolism | 12 | Glycine, serine and threonine metabolism | 6 |
| Alanine, aspartate and glutamate metabolism | 12 | Lysine degradation | 6 |
| RNA polymerase | 11 | Pantothenate and CoA biosynthesis | 4 |
| Fatty acid degradation | 11 | DNA replication | 7 |
| Fatty acid metabolism | 12 | Galactose metabolism | 4 |
| Arginine biosynthesis | 9 | N-Glycan biosynthesis | 5 |
| Endocytosis | 22 | Synthesis and degradation of ketone bodies | 3 |
| RNA transport | 17 | Phosphatidylinositol signaling system | 10 |
| alpha-Linolenic acid metabolism | 8 | Phosphonate and phosphinate metabolism | 3 |
| Glutathione metabolism | 9 | Tyrosine metabolism | 3 |
| Tryptophan metabolism | 7 | Valine, leucine and isoleucine biosynthesis | 2 |
| Fructose and mannose metabolism | 8 | One carbon pool by folate | 2 |

**Table S3.** Functional annotation of the 34 putative protein targets for BPA.

| Protein ID | Blastp (SwissProt) | Enzyme commission | Subcellular location (Deeploc) | InterPro (Protein Family) | PROSITE (signature) | CD-Search (ID) | KEGG (KO) | KEGG (pathways) |
|---|---|---|---|---|---|---|---|---|
| P00070270 | ABC transporter B family member 11 | 3.6.3.44 | Cell membrane (membrane) | Type I protein exporter (IPR039421) | PS50929 (ABC_TM1F); PS50929 (ABC_TM1F); PS50893 (ABC_TRANSPORTER_2); PS00211 (ABC_TRANSPORTER_1) | Multidrug resistance protein superfamily (cl36537) | K05658 | map02010; map04976; map05206; map05226 |
| P00300149 | Protein kinase dsk1 | - | Cytoplasm (soluble) | n.a. | PS50011 (PROTEIN_KINASE_DOM); PS00107 (PROTEIN_KINASE_ATP); PS00108 (PROTEIN_KINASE_ST) | Catalytic domain of the Serine/Threonine Kinase (cd14136) | K08832 | - |
| P00090254 | Phenylalanine-4-hydroxylase | 1.14.16.1 | Cytoplasm (soluble) | Aromatic amino acid hydroxylase (IPR001273) | PS51671 (ACT); PS51410 (BH4_AAA_HYDROXYL_2); PS00367 (BH4_AAA_HYDROXYL_1) | Biopterin-dependent aromatic amino acid hydroxylase superfamily (cl01244) | K00500 | map00360; map00400; map00790; map01100; map01230 |
| P00030265 | Casein kinase I isoform epsilon | 2.7.11.1 | Cytoplasm (soluble) | n.a. | PS50011 (PROTEIN_KINASE_DOM); PS00107 (PROTEIN_KINASE_ATP); PS00108 (PROTEIN_KINASE_ST) | Serine/Threonine protein kinase (Casein Kinase 1) (cd14016) | K08960 | map04011; map04068; map04310; map04340; map04341; map04390; map04391; map04392; map04710; map04711 |
| P00150104 | Extracellular signal-regulated kinase 1 | - | Cytoplasm (soluble) | n.a. | PS50011 (PROTEIN_KINASE_DOM); PS00107 (PROTEIN_KINASE_ATP); PS00108 (PROTEIN_KINASE_ST); PS01351 (MAPK) | Serine/Threonine Kinase (Mitogen-Activated Protein Kinase) (cd07834) | K04371 | - |
| P00210034 | Probable glutamine--tRNA ligase | 6.1.1.18 | Cytoplasm (soluble) | Glutamine-tRNA synthetase (IPR004514) | PS00178 (AA_TRNA_LIGASE_I) | Glutamine-tRNA ligase superfamily (cl31940) | K01886 | map00970; map01100 |
| P00210141 | Diphthine methyl ester synthase | 2.1.1.314 | Cytoplasm (soluble) | Diphthine synthase (IPR004551) | - | Diphthine synthase (PTZ00175) | K00586 | - |
| P00710081 | UDP-glucose 4-epimerase | 5.1.3.2 | Cytoplasm (soluble) | UDP-glucose 4-epimerase (IPR005886) | - | UDP-glucose 4 epimerase (subgroup 1) (cd05247) | K01784 | map00052; map00520; map01100 |
| P00100308 | Thymidylate kinase | 2.7.4.9 | Cytoplasm (soluble) | Thymidylate kinase (IPR018094) | PS01331 (THYMIDYLATE_KINASE) | Thymidylate kinase (cl17190) | K00943 | map00240; map01100 |
| P00420197 | Calmodulin | - | Cytoplasm (soluble) | n.a. | PS50222 (EF_HAND_2); PS00018 (EF_HAND_1) | Ca2+-binding protein (EF-hand) superfamily (cl34916) | K02183 | - |
| P00470113 | Extracellular signal-regulated kinase 1 | - | Cytoplasm (soluble) | Mitogen-activated protein (MAP) kinase; ERK3/4 (IPR008350) | PS50011 (PROTEIN_KINASE_DOM); PS00107 (PROTEIN_KINASE_ATP); PS00108 (PROTEIN_KINASE_ST) | Serine/Threonine Kinase (Mitogen-Activated Protein Kinase) (cd07834) | K04371 | - |
| P00040164 | Dual specificity protein phosphatase CDC14A | 3.1.3.16; 3.1.3.48 | Cytoplasm (soluble) | Tyrosine-protein phosphatase CDC14 (IPR026070) | PS50056 (TYR_PHOSPHATASE_2); PS00383 (TYR_PHOSPHATASE_1) | CDC14 family proteins (cd14499) | K06639 | map04110; map04111; map04113 |
| P00870013 | Geranylgeranyl pyrophosphate synthase penG | 2.5.1.1; 2.5.1.10; 2.5.1.29 | Cytoplasm (soluble) | Polyprenyl synthetase (IPR000092) | PS00444 (POLYPRENYL_SYNTHASE_2) | Polyprenyl synthetase (pfam00348) | K00804 | map00900; map01100; map01110; map01130 |
| P00450170 | ABC transporter E family member 2 | - | Cytoplasm (soluble) | RLI1 (IPR013283) | PS51379 (4FE4S_FER_2); PS00198 (4FE4S_FER_1); PS50893 (ABC_TRANSPORTER_2); PS00211 (ABC_TRANSPORTER_1) | Translation initiation factor RLI1 superfamily (cl34208) | K06174 | - |

| | | | | | | | | |
|---|---|---|---|---|---|---|---|---|
| P00340021 | Calcium-transporting ATPase 2 | 3.6.3.8 | Endoplasmic reticulum (membrane) | P-type ATPase (IPR001757) | PS00154 (ATPASE_E1_E2) | Haloacid Dehalogenase_like superfamily (cl21460) | K01537 | map04020; map04022; map04972; map05010 |
| P00220032 | Leucine aminopeptidase 2 | 3.3.2.6 | Endoplasmic reticulum (membrane) | Aminopeptidase; leukotriene A4 hydrolase-like (IPR034015) | PS00142 (ZINC_PROTEASE) | Peptidase M1 family (cd09599) | K01254 | map00590; map01100 |
| P00530165 | Calcium-transporting ATPase | 3.6.3.8 | Endoplasmic reticulum (membrane) | P-type ATPase (IPR001757) | PS00154 (ATPASE_E1_E2) | Haloacid Dehalogenase-like Hydrolases superfamily (cl21460) | K01537 | map04020; map04022; map04972; map05010 |
| P00300125 | 26S proteasome non-ATPase regulatory subunit 1 homolog B | - | Endoplasmic reticulum (soluble) | 26S Proteasome non-ATPase regulatory subunit 1 (IPR035266) | - | 26S proteasome regulatory complex component superfamily (cl34911) | K03032 | map03050; map05169 |
| P00430133 | Cathepsin D | 3.4.23.34; 3.4.23.5 | Lysosome (soluble) | Aspartic peptidase A1 family (IPR001461) | PS51257 (PROKAR_LIPOPROTEIN); PS51767 (PEPTIDASE_A1); PS00141 (ASP_PROTEASE) | Cellular and retroviral pepsin-like aspartate proteases superfamily (cl11403) | K01379 | map04071; map04140; map04142; map04210; map04915; map05152 |
| P00680119 | Ras-related protein Rab-8B | - | Golgi apparatus (membrane) | Small GTPase (IPR001806) | PS51419 (RAB); PS00675 (SIGMA54_INTERACT_1) | Rab subfamily of small GTPases (smart00175) | K07901 | map04144; map04152; map04530; map04972 |
| P00260129 | Ras-related protein Rab6 | - | Golgi apparatus (membrane) | Small GTPase (IPR001806) | PS51419 (RAB) | Rab GTPase family 6 (Rab6) (cd01861) | K07893 | - |
| P00700035 | Ras-related protein ORAB-1 | - | Golgi apparatus (membrane) | Small GTPase (IPR001806) | PS51419 (RAB) | Rab GTPase families 8, 10, 13 (Rab8, Rab10, Rab13) (cd01867) | K07874 | map05134 |
| P00680028 | Phenylalanine-4-hydroxylase | 1.14.16.1 | Mitochondrion (soluble) | Aromatic amino acid hydroxylase (IPR001273) | PS51671 (ACT); PS51410 (BH4_AAA_HYDROXYL_2); PS00367 (BH4_AAA_HYDROXYL_1) | Biopterin-dependent aromatic amino acid hydroxylase superfamily (cl01244) | K00500 | map00360; map00400; map00790; map01100; map01230 |
| P00530125 | Peptidyl-prolyl cis-trans isomerase CYP19-2 | 5.2.1.8 | Mitochondrion (soluble) | n.a. | PS50072 (CSA_PPIASE_2); PS00170 (CSA_PPIASE_1) | Cyclophilin superfamily (cl00197) | K01802 | - |
| P00620128 | Phenylalanine--tRNA ligase | 6.1.1.20 | Mitochondrion (soluble) | Phenylalanyl-tRNA synthetase; class IIc; mitochondrial (IPR004530) | PS50862 (AA_TRNA_LIGASE_II); PS51447 (FDX_ACB) | Phenylalanine-tRNA synthetase superfamily (cl33567) | K01889 | map00970 |
| P00800061 | Probable succinyl-CoA:3-ketoacid coenzyme A transferase | 2.8.3.5 | Mitochondrion (soluble) | n.a. | - | Sugar Phosphate Isomerase superfamily (cl00339) | K01027 | map00072; map00280; map00650 |
| P00930049 | Guanylate kinase 1 | 2.7.4.8 | Mitochondrion (soluble) | Guanylate kinase (IPR017665) | PS50052 (GUANYLATE_KINASE_2); PS00856 (GUANYLATE_KINASE_1) | Guanylate kinase (TIGR03263) | K00942 | map00230; map01100 |
| P00370082 | DNA replication licensing factor mcm7 | 3.6.4.12 | Nucleus (soluble) | DNA replication licensing factor Mcm7 (IPR008050) | PS50051 (MCM_2); PS00847 (MCM_1) | Minichromosome maintenance proteins (MCM) superfamily (cl33383) | K02210 | map03030; map04110; map04111; map04113 |
| P00950041 | Serine/threonine-protein kinase AFC3 | 2.7.12.1 | Nucleus (soluble) | n.a. | PS50011 (PROTEIN_KINASE_DOM); PS00108 (PROTEIN_KINASE_ST) | Protein Kinases_like superfamily (cl21453) | K08287 | - |
| P00040322 | Dual specificity protein kinase CLK4 | - | Nucleus (soluble) | n.a. | PS50011 (PROTEIN_KINASE_DOM); PS00108 (PROTEIN_KINASE_ST) | Protein Kinases_like superfamily (cl21453) | K23561 | - |

| | | | | | | | | |
|---|---|---|---|---|---|---|---|---|
| P00150260 | Cyclin-dependent kinase 10 | 2.7.11.22 | Nucleus (soluble) | n.a. | PS50011 (PROTEIN_KINASE_DOM); PS00107 (PROTEIN_KINASE_ATP); PS00108 (PROTEIN_KINASE_ST) | Cyclin-Dependent protein Kinase-like Serine/Threonine Kinases (cd07829) | K02449 | - |
| P00300132 | DNA-directed RNA polymerase I subunit rpa1 | 2.7.7.6 | Nucleus (soluble) | DNA-directed RNA pol I; largest subunit (IPR015699) | PS00253 (INTERLEUKIN_1) | Largest subunit (RPA1) of eukaryotic RNA polymerase I (RNAP I) (cd01435) | K02999 | map00230; map00240; map01100; map03020 |
| P00100039 | Peroxisomal acyl-coenzyme A oxidase 1 | - | Peroxisome (membrane) | Acyl-CoA oxidase (IPR012258) | - | Acyl-CoA dehydrogenase superfamily (cl33189) | K00232 | - |
| P00070322 | Alkyl-DHAP synthase | 2.5.1.26 | Peroxisome (soluble) | Alkyl dihydroxy acetonephosphate synthase (IPR025650) | PS51387 (FAD_PCMH) | FAD/FMN-containing dehydrogenase (COG0277) | K00803 | map00565; map01100; map04146 |

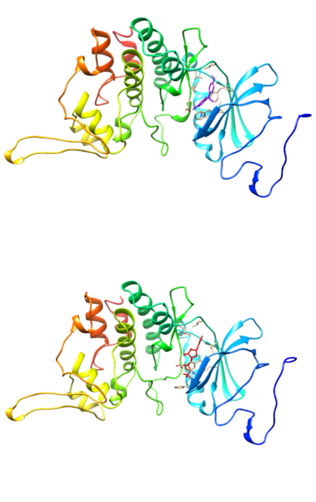

P00950041

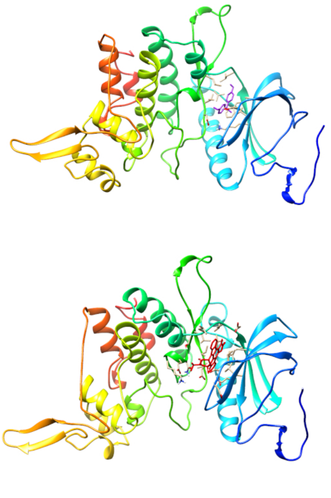

P00040322

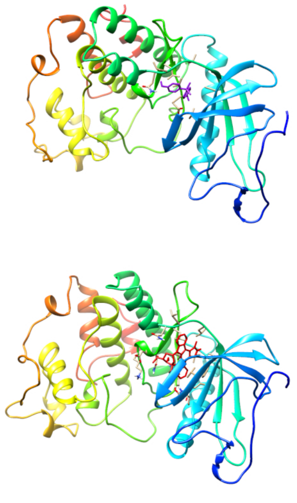

P00300149

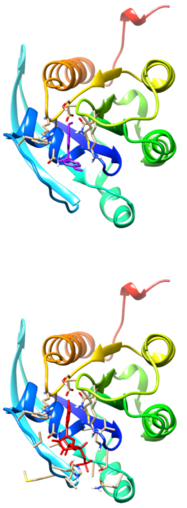

P00680028

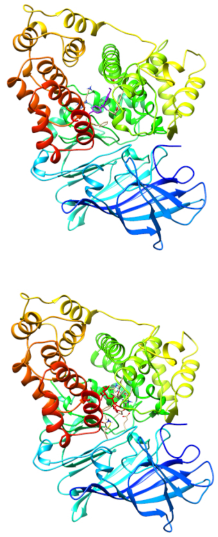

P00220032

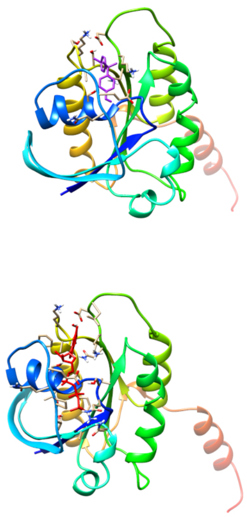

P00260129

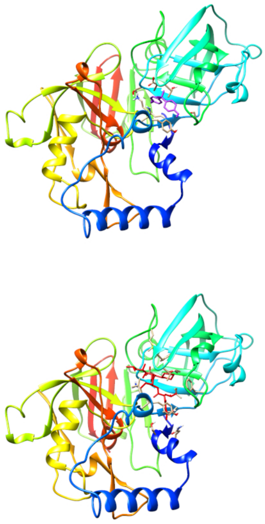

P00430133

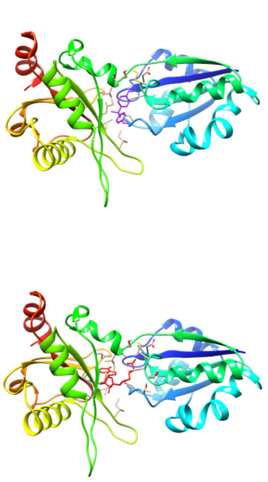

P00210141

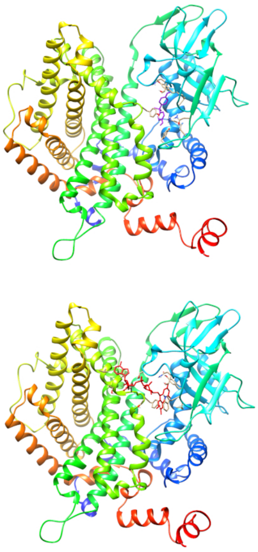

P00100039

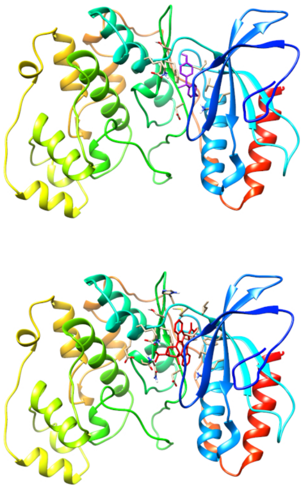

P00150104

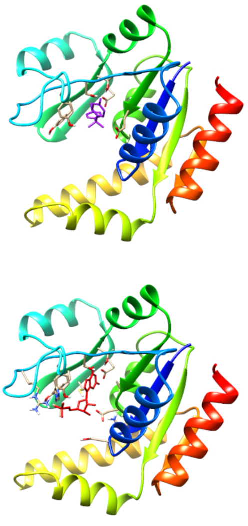

P00930049

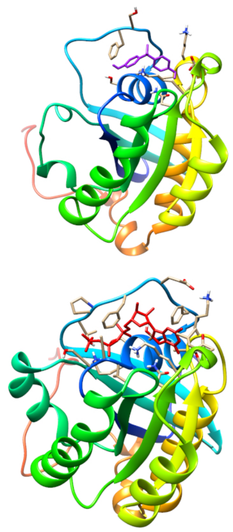

P00700035

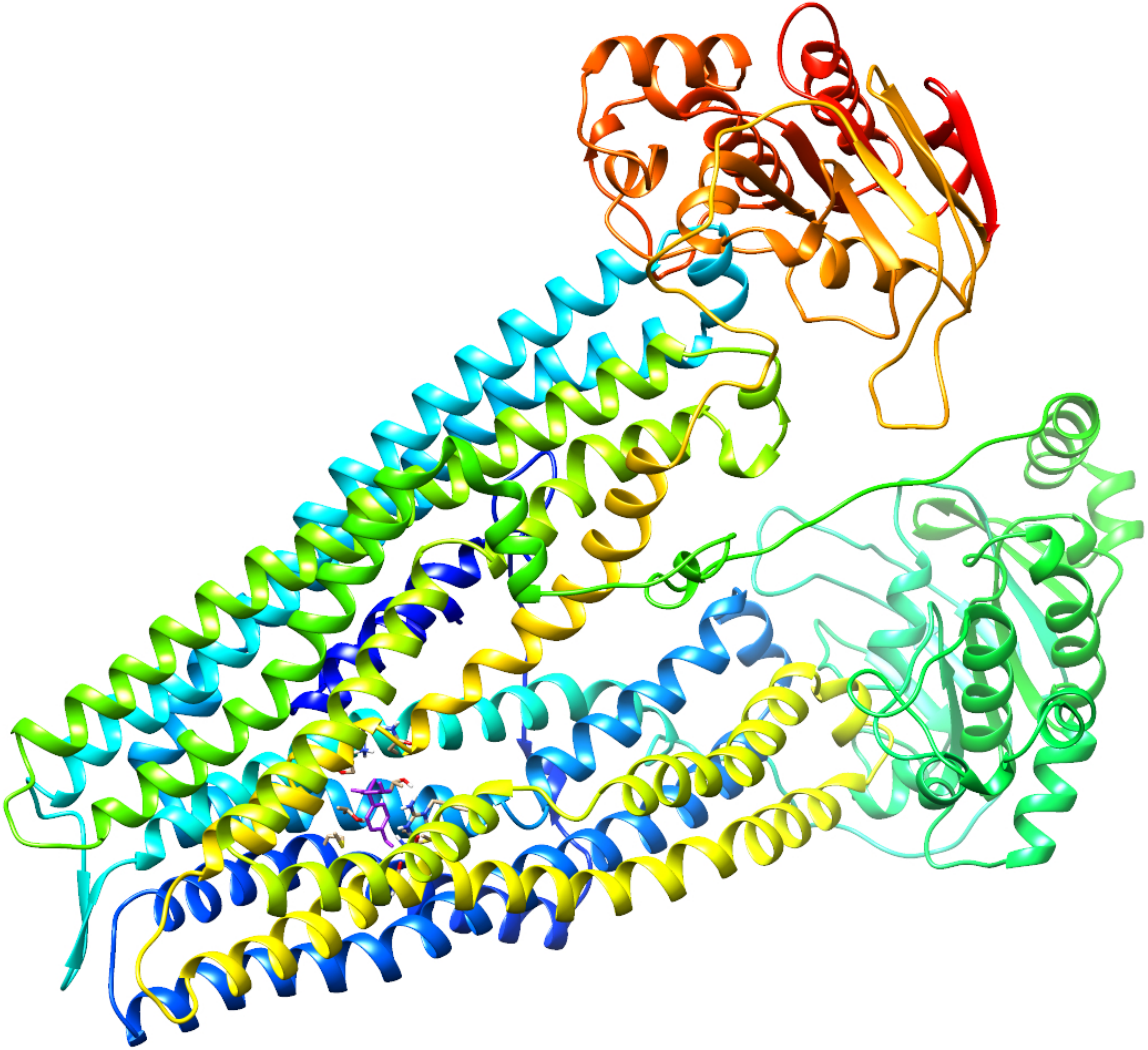

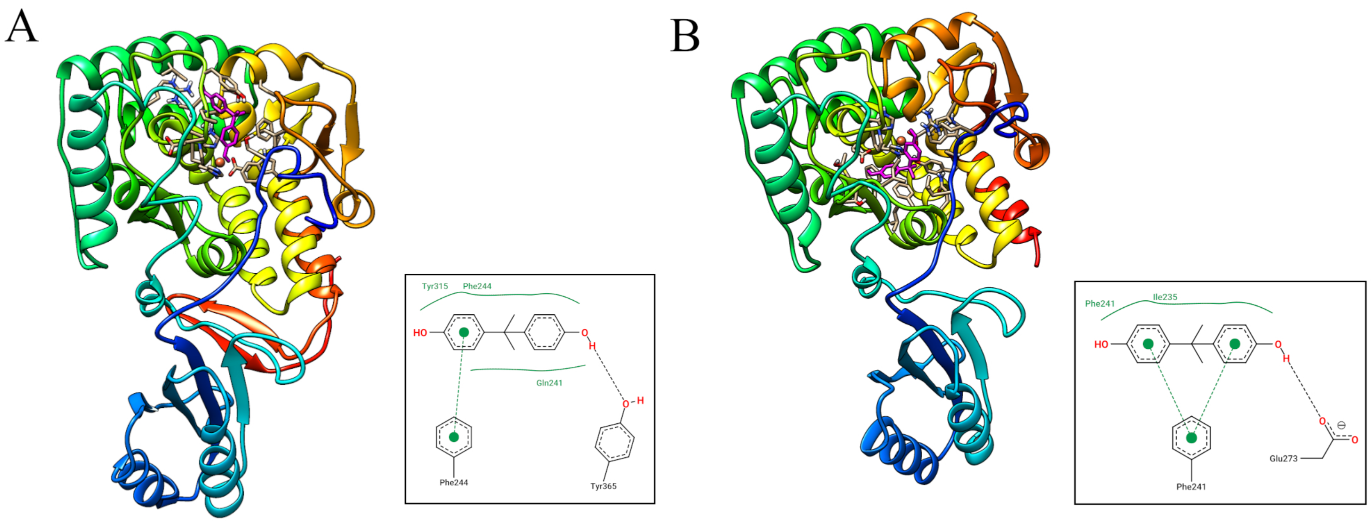
A
Tyr315
Phe244
HO
O
H
Gln241
O
H
Phe244
Tyr365
B
Phe241
Ile235
HO
O
H
O
O
Glu273
Phe241
